# Targeted Analysis of >1500 Plasma Proteoforms via Individual Ion Mass Spectrometry

**DOI:** 10.64898/2026.05.28.728622

**Authors:** Aniel Sanchez, Indira Pla, Katrina N. Peterson, Vincent White, Troy D Fisher, Michael A. R. Hollas, Taojunfeng Su, Nhat Hoang Van Le, Claire R. Harrington, Paolo Cravedi, David N. Assis, Paola Barrios, Therese Elaine Banea, Daniela P. Ladner, Eleonora Forte, Matthew Lucky, John T. Wilkins, Douglas E Vaughan, Michael A. Caldwell, John P. McGee, Neil L. Kelleher

**Author notes:** Corresponding Author: Neil L. Kelleher.

## Abstract

Plasma proteomics has long sought to accessibly sense human biology, precisely detect disease states, and advance diagnostics through clinical translation. In recent years, companies with defined panels such as Olink and Somascan have entered the field to complement bottom-up mass spectrometry (BUP). This study leverages a novel mass spectrometry platform to capture targeted proteoform information lost by mainline antibody-, aptamer-, and BUP-driven workflows. The Plasma Proteoform Assay (PPA) uses Individual Ion Mass Spectrometry (I²MS) to resolve mixtures of intact proteins presented by direct injection. Two proteoform panels, **PPA 526** and **PPA 1514,** were defined from human plasma samples obtained from 81 individuals. The panels quantify 526 proteoforms derived from 59 genes and 1,514 proteoforms from 155 genes, respectively. Reproducibility for both panels showed coefficients of variation below 20% for most proteoforms (59-80%). PPA was benchmarked in studies including subjects with hepatic cirrhosis (N=30) and resilient agers carrying a SERPINE1 (PAI 1) mutation (N=27). PPA signatures distinguished SERPINE1 mutation carriers from affected individuals and provided sufficient resolution to discriminate among cirrhosis disease stages. In summary, we present **PPA 526** and **PPA 1514**, the first scalable plasma proteoform panels capable of tracking hundreds to thousands of targets in a few minutes per sample.

## Introduction

Plasma, the liquid fraction of blood, retains most of the physiological properties of whole blood and is widely used in clinical applications due to its minimally invasive collection ^1, 2^. However, comprehensive characterization of the plasma proteome remains challenging, particularly for proteins that occur in multiple forms, now called “proteoforms”^3^. This is in part due to the extreme dynamic range of protein concentrations (spanning more than 10 orders of magnitude) and the overwhelming presence of a few highly abundant proteins (*e.g.*, albumin and immunoglobulins, which together account for ∼99% of total plasma protein content). For proteoforms specifically, their molecular uniqueness from one another means that measuring them is limited by their individual abundances, which can be an order of magnitude or lower than the corresponding protein abundance. Thus, the complexity of the plasma proteome poses significant analytical hurdles to routinely sensing human biology.

The proteomics field has responded to the hurdles of plasma proteomics by largely forgoing proteoform-level biomarker detection and developing both advanced affinity-based readouts and bottom-up proteomics (BUP) workflows. One such affinity-based technology is Olink’s Proximity Extension Assay (PEA), which has scaled in both cohort size and proteome depth in recent years. Early studies using PEA profiled approximately 1,500 individuals for 233 proteins ^4^, while mid-scale efforts such as the China Kadoorie Biobank analyzed ∼2,000 individuals for 2,923 proteins ^5^. This was followed by the SCALLOP consortium, which examined ∼31,000 individuals for ∼90 cardiovascular proteins^6^. Most recently, the UK Biobank Pharma Proteomics Project has expanded to >50,000 participants with ∼3,000 proteins using Olink Explore panels ^7^, marking a major leap in scalable plasma proteomics. However, large comparative studies have noted limited consensus among next-generation immunoaffinity assays, presumably due to the lack of proteoform-level information ^8, 9^. Using BUP, the largest cohort studies typically analyze hundreds to >1000 individuals, achieving depths of hundreds to a few thousand proteins per sample when using high-throughput plasma workflows optimized for robustness and reproducibility ^10–12^. Very recently, Vincent *et al.*^13^ reported a technically deeper plasma proteomics workflow with potential for ∼100 samples per day with a depth of >4000 proteins.

Mass spectrometry-based analysis of plasma and serum analyses have predominantly relied on BUP ^14–17^. This approach identifies proteins through their enzymatically generated peptides; however, BUP loses critical information about the original proteoforms by confounding similar peptides from distinct proteoforms^3^. To capture more human biology, Top-Down Proteomics (TDP) presents an alternative strategy, enabling direct analysis of intact proteins from plasma or serum using discovery-driven or targeted workflows ^18–23^. Despite these advances, current implementations of TDP often require extensive fractionation, including liquid chromatography and capillary electrophoresis, which limits scalability for high-throughput or clinical applications due to high complexity and cycle time^24^. We previously demonstrated that Top-Down analysis of soluble human plasma proteoforms via acetonitrile precipitation is reproducible and capable of capturing biologically relevant patterns, as evidenced by our LC-MS analysis of a liver cirrhosis cohort (n = 30 subjects)^23^. The missing piece, however, was the ability to compete with the throughput of other novel proteomics workflows at a clinical scale.

In this study, we introduce a direct plasma proteoform analysis workflow leveraging detection of individual ions (I²MS)^25^ in an optimized configuration integrating mass spectrometry with Field Asymmetric Ion Mobility Spectrometry (FAIMS)^26^ and a Sample Stream platform ^27^. Using the acetonitrile-soluble plasma fraction and a deep knowledge of proteoforms ^23^ arranged into a reference atlas ^28^, we achieved the scalable monitoring of more than 1,500 proteoforms across a cohort of >100 subjects in as little as 10 minutes per subject. We also benchmarked basic performance specifications of defined panels, **PPA 526** and **PPA 1514**, to sense the biology encoded in a readily accessible set of proteoforms in the human plasma proteome.

## Methods

### Plasma Samples

To test the generality of the PPA panels developed in this study, we analyzed plasma samples spanning a range of disease phenotypes, including conditions known to perturb abundant plasma proteoforms. Samples were organized into three analytical sets (Table 1), each serving a distinct purpose**. Set 1** (N = 81) was used for panel discovery and qualitative proteoform definition. This set included plasma samples obtained from STEMCELL Technologies (Vancouver, BC, Canada; N = 1 healthy reference) and from the Comprehensive Transplant Center at Northwestern Medicine, comprising cohorts of individuals, including patients with liver cirrhosis (N = 40, including 10 controls) ^23^, patients with autoimmune hepatitis (AIH; N = 40, including 10 controls).

**Table 1.** Study cohorts used for PPA development and benchmarking.

| Study Set | Cohort description |
| --- | --- |
| Set 1 | AIH (N=40) *, Cirrhosis—Discovery (N=40), Healthy reference (N=1) |
| Set 2 | SERPINE1 (N=27) |
| Set 3 | Cirrhosis—Stage stratified (N=30) |

\*AIH, Autoimmune Hepatitis

**Set 2** and **Set 3** were used for streamlined quantitative analysis to assess condition dependent differences in proteoform abundance. **Set 2** comprised an independent cohort of individuals from the Human Longevity Laboratory (HLL; N = 27). **Set 3** consisted of a previously evaluated cirrhosis cohort representing three clinical stages (N = 30) ^23^, comprising 9 compensated, 10 compensated with portal hypertension, and 11 decompensated individuals, which were also included in **Set 1** for panel definition. As in the original analysis, samples were evaluated in pooled form by disease stage (three pools per condition), while the individual level composition of the cohort is reported here to describe its clinical structure ^23^.

Typically, 50 microliters were used for sample preparation and 1/10^th^ of the resultant samples were used for technical replicate injections onto the SampleStream device. These studies adhered to NIH guidelines for human subject research. The Northwestern IRB granted a Waiver of Consent and approved the study under protocols STU00216399 and STU00217037 (cirrhosis), and STU00216136 for the HLL.

### Execution of proteoform panels (PPA 526 and PPA 1514)

Plasma samples from controls and patients with liver cirrhosis or autoimmune hepatitis (**Set 1**, N = 81) were processed using a previously published acetonitrile precipitation protocol called SPAP ^23^. Proteoforms from the resulting supernatants were cleaned via SampleStream (SS) and directly injected into an Orbitrap mass spectrometer capable of individual ion MS (I^2^MS, commercialized as Direct Mass Technology)^29^, either without or with FAIMS (see **Figure 1**).

**Figure 1.**
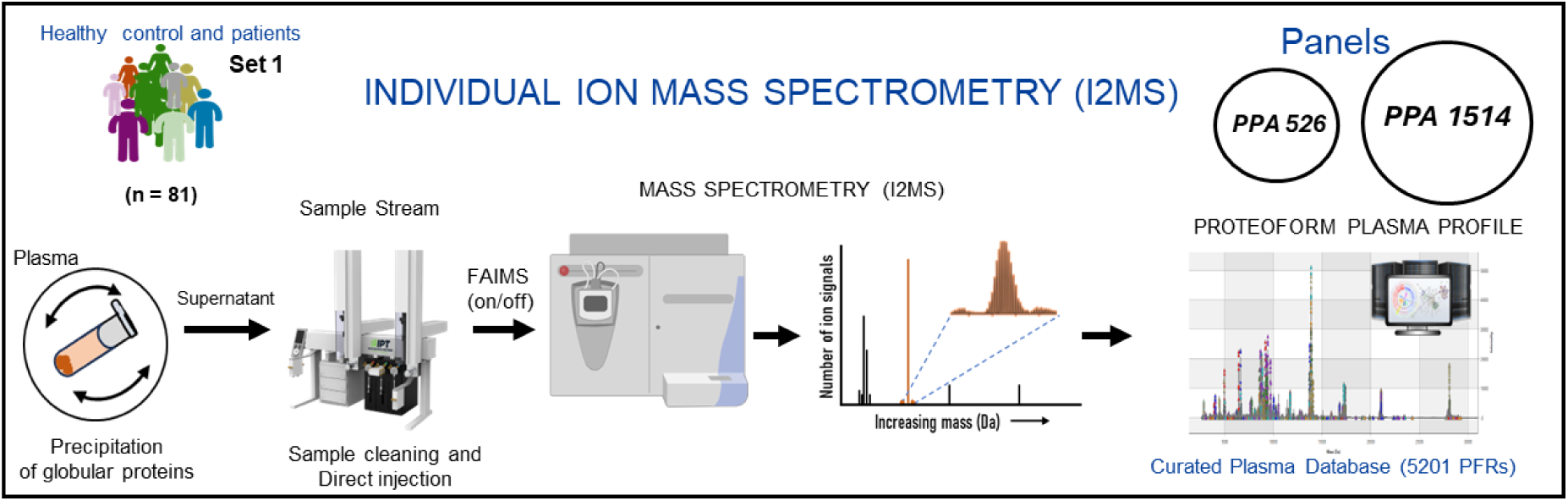
Execution of proteoform panels using direct injection and individual-ion mass spectrometry (I²MS). Through iterative analysis of plasma samples from **Set 1**, comprising 81 individuals, a healthy control and patients with liver diseases, we defined two plasma proteoform panels: **PPA 526** and **PPA 1514** (upper right). From left to right: A rapid protocol was applied to deplete high-abundance proteins, retaining a fraction of proteoforms in solution (SPAP fraction) for top-down mass spectrometry. Proteoforms were injected into the mass spectrometer either with or without FAIMS, after sample cleaning using a SampleStream device, and each proteoform was measured by I²MS. A bioinformatic pipeline incorporating a reference set of >5000 known proteoforms from MS² experiments enables MS¹ feature detection, proteoform identification and quantification.

Proteoforms were identified through feature detection in mass space using a curated Plasma Proteoform Repository, annotated from extensive prior LC–MS/MS analyses ^23, 27, 30^ performed on the same or related samples (including most of those analyzed here), as well as I²MS² data. From these efforts, we assembled a set of 5,201 high quality reference plasma proteoforms and used these experimentally verified species as the basis for their targeted detection and quantification (**Figure 1**, right). Based on targeted proteoform detection of such library proteoforms, we defined two panels that either use FAIMS (**PPA 1514**) or do not (**PPA 526**) and assessed their performance characteristics.

### Single shot plasma proteoform preparation (PPA 232)

Plasma samples were prepared for direct single-shot analysis without acetonitrile precipitation. Briefly, plasma was diluted to 0.14% v/v using 0.09% v/v formic acid in water. The final dilution was used directly for I^2^MS analysis, using FAIMS. **PPA 232** enabled us to report basic comparisons of observable proteoforms and MS acquisition times under matched instrumental conditions for **PPA 232** relative to **PPA 1514**, with the latter panel using the extra step of ACN precipitation.

### Chemicals and reagents

Acetonitrile (ACN), Water LC-MS grade, and formic acid were purchased from Thermo Fisher Scientific.

### Extraction and Recovery of Proteoforms from Human Plasma

Acetonitrile (ACN) precipitations from plasma were performed, followed by a protocol previously published (**Figure 1**, left) ^23^. Briefly, 50 or 100 µL of plasma was centrifuged at 13,000 rpm for 30 minutes at room temperature to separate the lipid layer. After discarding the lipid layer and optionally incorporating 0.2% formic acid, an equal volume of cold acetonitrile (ACN), stored at −20°C, was added to the sample, followed by vortexing for 15 seconds. Samples were incubated on ice, clarified via centrifugation and precipitate removal, and dried or aliquoted before reconstitution in 0.1% formic acid for analysis. During the optimization process, protein concentration was measured using a BCA assay (Thermo Fisher Scientific) according to the manufacturer’s instructions.

### Data Acquisition by Individual Ion Mass Spectrometry

I²MS analysis was performed on either a modified Q Exactive HF Hybrid Quadrupole-Orbitrap Mass Spectrometer ^31^ or an Orbitrap Exploris 480, both coupled to the SampleStream device for sample clean up, concentration and injection (**Figure 1**, middle). Using SampleStream for buffer exchange above a 5 kDa molecular weight cutoff membrane, a denaturing buffer was introduced (0.2% formic acid and 30% acetonitrile in Optima-grade water). After this cleanup and sample concentration, a range of 22 ng to 1 µg of total sample was directly infused into the mass spectrometer at a 1.6 µL/minute flow rate via a heated electrospray ionization (HESI) source. Injection technical replicates were performed at least in duplicate, with some experiments including up to 5–10 replicates.

For experiments using FAIMS coupled to the Orbitrap Exploris 480, HESI parameters included a spray voltage of 3.5–3.9 kV and a capillary temperature of 360°C. Nitrogen flow was maintained at 4.6 L/minute under standard resolution settings. Multiple compensation voltages were evaluated to identify conditions that best reduced APOA1, a highly abundant protein in these fractions^23^ (**Figure S3**). A compensation voltage of −35 V was used for most experiments (from 1 to 40 minutes acquisitions), whereas others employed an equal split between −35 V and −40 V. All experiments employed a fixed injection time of 10 ms. Additional settings included a resolving power of 240,000 at 200 m/z, 1 µscan, and a trapping gas pressure of 0.3 (arb).

Without FAIMS, the ion acquisition required 47 minutes using AIC ^32^. When applicable, samples were analyzed in a randomized order of biological replicates to minimize batch effects across treatments.

### Data Processing

Data acquired as I^2^MS STORI (Selective Temporal Overview of Resonant Ions)^33^ files were compressed and submitted to a custom I^2^MS processing pipeline on the Quest high-performance computing cluster at Northwestern. This pipeline applies STORI analysis followed by iterative voting charge assignment to extract individual-ion charge information, allowing the reconstruction of mass-domain spectra ^25,33^. Subsequently, the resulting .dmt (direct mass technology) files were calibrated using an in-house Python workflow that applies an internal mass calibration based on known proteoform masses, correcting systematic mass offsets. Mass calibrated .dmt files were reprocessed in the I^2^MS data processing pipeline for grouped-run voting. Two separate merging techniques were attempted: 1) .dmt files of the same treatment were combined and voted together, and 2) all .dmt files from the study were combined and voted together. These merged .dmt files were then ‘split’ using a Python script to separate ions according to each acquisition. An overview of this processing workflow is detailed in **Figure S1**.

### Processing of I^2^MS Data

A proteoform reference database was generated as a modified FASTA file containing ProForma^34^ sequence information. The database was constructed using extensive LC–MS/MS plus I²MS² analyses of samples from the present study and from closely related cohorts^23, 27, 30^. Proteoform reference entries derived from LC-MS/MS data were curated by filtering at a conservative false discovery rate (FDR) of <1%^35^, resulting in a high confidence set of 5,201 plasma proteoforms (**Figure S1A**). The curated set of targeted proteoforms was stored in the FASTA file and used for proteoform identification in sample **Set 1** (**Table 1**), where .dmt files were searched using an in-house C# script to perform intact mass tag (IMT)^36^ matching against the set of known proteoforms. Maximum and sub tolerances were set to 10 and 5 ppm, respectively.

**PPA 526** and **PPA 1514** were defined using PFR identifications derived from **Set 1**, while **PPA 232** was defined from **Set 2** together with the reference healthy plasma sample. For **Sets 2** and **3**, as well as the reference sample, proteoforms corresponding to **PPA 526**, **PPA 1514**, or **PPA 232** were subsequently targeted and quantified using a single ion-based quantification approach. Briefly, ions within theoretical *m/z* windows for all possible charge states for a given proteoform are counted and summed, yielding a proteoform ion count for a given .dmt file. To achieve robust quantification supported by reliable precursor-level evidence, proteoform features within a single .dmt file were interrogated using IMT matching to clear isotopic distributions in MS1 (*i.e*., those with good MS1 evidence). The distribution of the number of quantified ions (log₂-transformed) was compared between proteoforms detected with and without good MS1 evidence (**Figure S1B**). We implemented a lower threshold for ion counts set to include 95% of ions consistent with the mass value of a targeted proteoform. Only proteoforms quantified with an ion count exceeding this cohort-specific threshold were retained for downstream analyses.

### Data Quality Control and Differential Expression

Variability introduced during sample processing was evaluated by calculating, for each proteoform, the coefficient of variation (CV) across technical and injection replicates using intensity values normalized to total ion count for each .dmt file. Additionally, the covariance was calculated. For each proteoform, we fit a random intercept mixed model with random effects for biological condition, biological replicate, and technical replicate (*PFRabundance ∼ −1 + (1|BioCond) + (1|BioRep) + (1|TechRep)).* For this analysis, intensities were log_2_-transformed and z-score normalized across runs. Variance components were extracted from the fitted model using the ‘lme4::VarCorr’ ^37^ function in R, and the percentage contribution of each component was computed as 100×vcov/∑vcov. Regression analysis, density curve graphs, and dynamic range plots were generated using the ‘ggplot2’ package (version 4.0.0) in R ^38^. Venn diagrams were generated using the BioVenn web application (https://www.biovenn.nl/) ^39^, to compare datasets according to proteoform identification numbers. All remaining figures were generated using GraphPad Prism (version 10.2.3).

Furthermore, identified proteins were matched to the Human Protein Atlas database (https://www.proteinatlas.org/) using SwissProt human accession numbers to facilitate grouping by protein abundance and comparison with publicly available protein data from the Olink Explore 3072 platform ^40^. The distribution of post-translational modifications (PTMs) present in each of the panels was shown through bar charts (‘ggplot2’ R package, version 4.0.0) and Sanky plots (‘ggsankey’, version 0.0.99999, R package) to show the association between biological processes and PTMs per proteoform family. Significant biological processes were accessed after doing gene ontology (GO) enrichment analyses using the STRING database (https://string-db.org/) ^41^ and the ‘clusterProfiler’ (version 4.10.1, R package) ^42^.

To detect proteoforms differentially expressed between biological conditions (*i.e.*, groups), we filtered the data to work with proteoforms quantified in at least 50% of the samples within a group. The analysis was done in RStudio (v 2026.01.1) ^43^ using the z-score normalized data described above. A mixed-effects model was fit for hierarchical evaluation of fixed (i.e., groups) and random factors (e.g., BioRep and TechRep) using the R package ‘lme4’ (version 1.1.37) ^37^. P-values were adjusted for multiple testing using the method of Benjamini–Hochberg ^44^, and proteoforms with adjusted p < 0.05 were considered differentially expressed. The result of these steps is a spreadsheet containing a quantitative output of expression for targeted proteoforms in each PPA panel and a metric of statistical confidence for each observation. The ability of differentially expressed proteoforms (DEPs) to discriminate between biological conditions was assessed by performing principal component analysis (PCA) and hierarchical clustering combined with heatmap visualization. The PCA analysis was done using the ‘stats::prcomp’ R function and visualized through the ‘fviz_pca_ind’ function of the ‘factoextra’ (version 1.0.7, R package). Heatmaps were generated with the z-scored normalized intensities of differentially expressed proteoforms (DEPs) using the R package ‘ComplexHeatmap’ (version 2.18.0) ^45^.

## Results and Discussion

### Proteoforms Panels: Expanding Dynamic Range of Plasma Proteoforms via FAIMS

**PPA 526** and **PPA 1514** were established and refined from plasma analysis of many individuals as described above and derive from 59 or 155 human genes as shown in Figure 2A and listed in **Tables S1** and **S2**, respectively. Surprisingly, despite containing 2.6-fold more proteoforms, PPA 1514 can be analyzed using FAIMS technology in less overall time (10–20 min with FAIMS vs 47 min), representing a two- to fourfold reduction.

**Figure 2.**
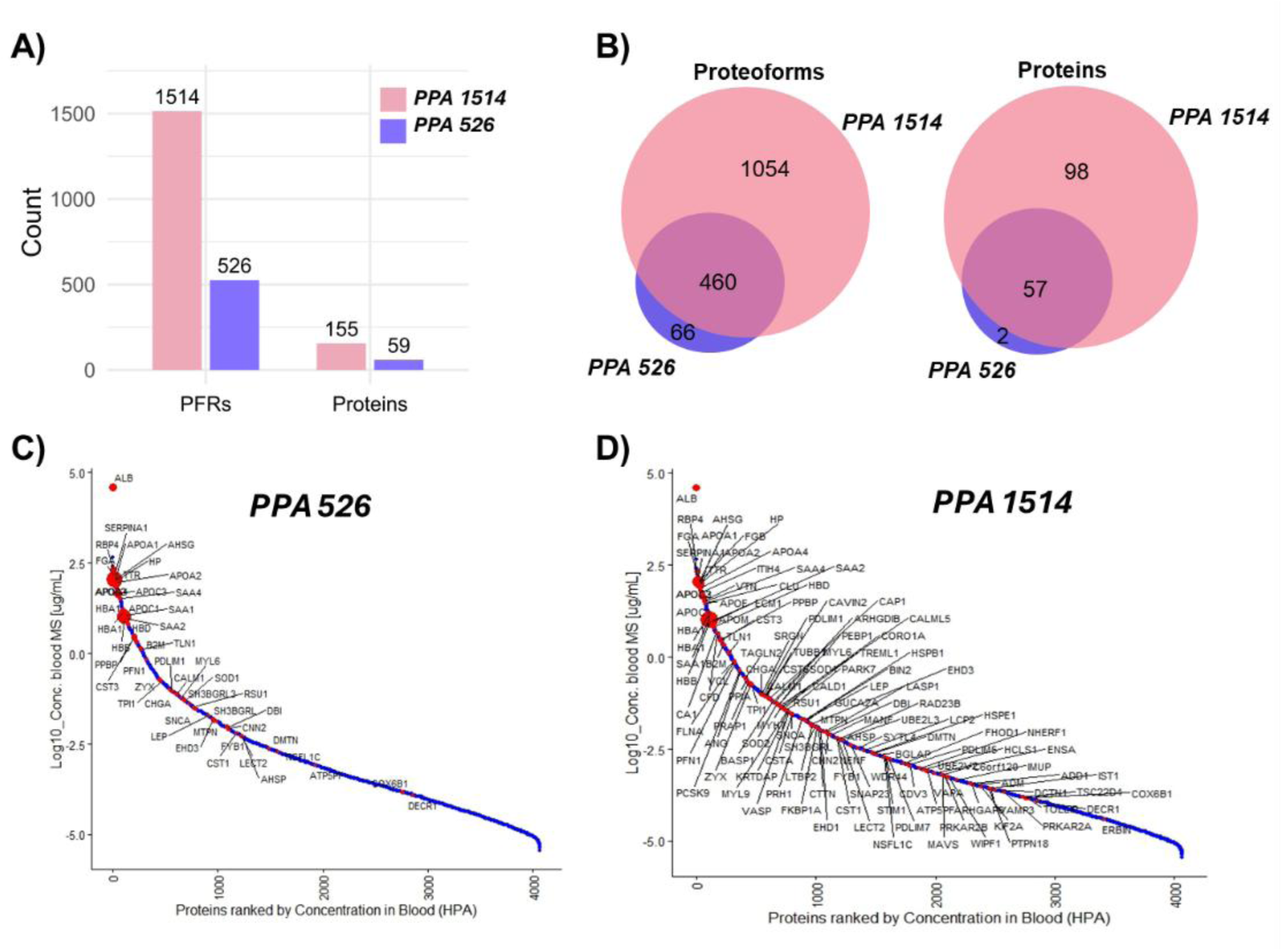
General features of proteoform (PFR) quantification in the two PPA Panels. **A)** Bar plots showing the number of PFRs and proteins identified in **PPA 526** (FAIMS off) and **PPA 1514** (FAIMS on). **B)** Venn diagram comparing PFR and protein identifications between **PPA 526** and **PPA 1514. C)** and **D)** Waterfall plots illustrate the dynamic range of proteins identified without FAIMS (**PPA 526**) and with FAIMS (**PPA 1514**), respectively. Protein ranks are based on expression levels reported in the Human Protein Atlas (red points; larger circles indicate more PFRs per protein), compared to HPA annotations for over 4,000 proteins (blue points). Notably, the number of unique proteins identified increased by 2.6-fold, indicating that the higher PFR counts were broadly distributed across protein families rather than driven solely by a few high-abundance families enriched through FAIMS (**PPA 1514**). The mass range of detected proteoforms remained similar to both panels, with both panels exhibiting bimodal distributions of PFR mass values, dominated by proteoforms <25 kDa (**Figure S2**).

More than 87% (460 out of 526) of the proteoforms identified in **PPA 526** were also detected in **PPA 1514** (**Figure 2B**). Similarly, the number of unique proteins confirmed was even higher, with 57 out of 59 (97%) proteins sampled in **PPA 526** also observed in **PPA 1514**. This high overlap was not unexpected, as we mostly selected a compensation voltage of –35 V to maintain the stability of most MS signals while simultaneously depleting APOA1, the most intense signal, during ion mobility separation (**Figure S2**).

In general, the 2.6-fold increase in proteins identified after FAIMS represented a subset of proteins that were clearly less abundant compared to those detected without additional gas phase fractionation. The proteoforms identified following FAIMS analysis were more densely distributed among proteins ranked beyond 1,000 in abundance (corresponding to concentrations around—or below—10 ng/mL; see **Figures 2C, D**). This concentration range is often associated with blood biomarker candidates ^17, 46^ and is commonly used as a threshold for classifying medium to low-abundance proteins.

The overall mass distribution of proteoforms identified in both panels was similar; however, we observed a slight increase in lower mass proteoforms in the **PPA 1514** (**Figure S3A)**. Examination of modifications revealed that many proteoforms in this lower-mass region corresponded to truncated proteins (**Figure S3B**). Truncated proteoforms in plasma/serum have been frequently reported^17, 23, 24, 47^. Formally, the biological relevance of each truncation can be very high for functional variants or low from artifacts of sample handling; this open question can be resolved in time by observing functional proteoforms repeatedly across the human subjects by deploying PPA panels at population scale. Notably, we previously demonstrated that certain truncated forms can exhibit drastic changes between biological conditions ^23, 48^, particularly in plasma, where protein-binding interactions are essential ^49^ and many pro-enzymes are activated through programmed proteolytic clipping. Additionally, these panels contain proteoforms with phosphorylation, acylation, disulfide bonds, and a variety of other PTMs (**Figure S3B**).

### Functional Categories of Protein Expression into Proteoforms

The **PPA 526** and **PPA 1514** panels contain a total of 1,580 proteoforms originating from 157 genes (**Tables S1, S2**). Together, these constitute the first version of our plasma proteoform reference set accessible by scalable I²MS analysis. Proteoforms identified in **PPA 526** vs. **PPA 1514** largely correspond to genes whose corresponding proteins (59 vs. 155, respectively) ranked among the top 3,000 most abundant plasma proteins in the Human Protein Atlas (HPA) database (**Figure 2C, D**). This observation is consistent with the expected abundance-driven detectability of intact proteoforms in clinical plasma samples.

Both **PPA 526** and **PPA 1514** show three recurring biological themes (**Figure 3**): a lipid/cholesterol transport module, a cytoskeleton/actin module, and a defense–platelet/acute phase module. In the lipid theme, GO terms such as reverse cholesterol transport, cholesterol efflux/clearance, and HDL particle remodeling/assembly align strongly with apolipoproteins (APOA1, APOA2, APOA4, APOC1–3, APOM) and PCSK9, covering a large subset of proteins related to lipid transport and metabolism.

**Figure 3.**
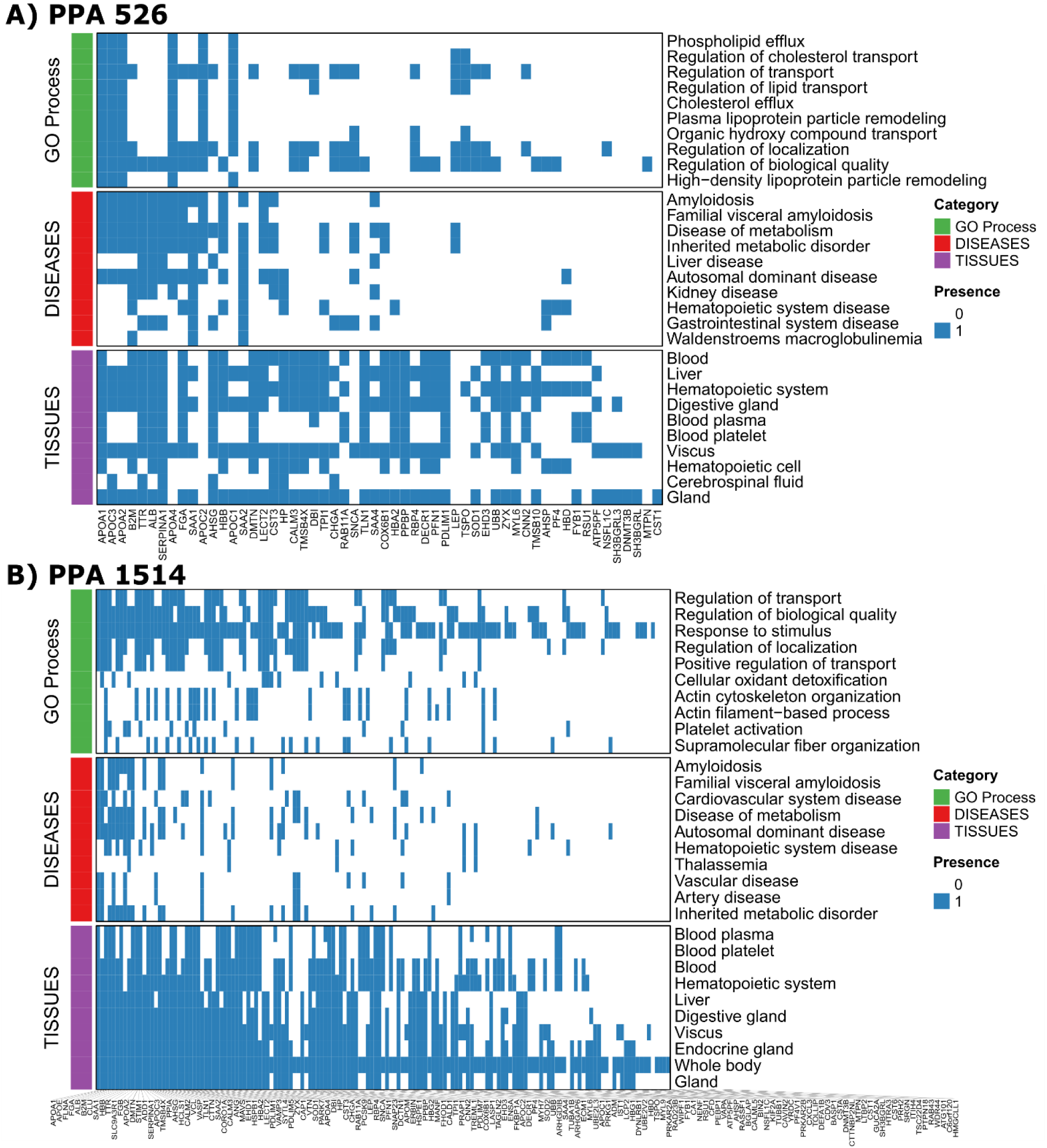
Protein membership across enriched Gene Ontology processes, organs/tissues, and disease categories is shown as binary heatmaps. **A)** The **PPA 526** Panel is annotated to show the distribution across individual 59 proteins, preserving the full structure of term–gene relationships across functional, tissue, and disease dimensions. **B)** The **PPA 1514** Panel presents the same annotations after reordering proteins to group shared memberships, yielding a more compact representation that facilitates comparison across molecular processes, organs/tissues, and disease categories. Categories are indicated by colors (right).

At the organ/tissue level, the lipid-associated module in **Figure 3** maps most clearly to liver and blood/plasma tissue annotations, consistent with enrichment for lipid and lipoprotein-related transport processes. Disease annotations associated with this block fall into metabolic and cardiovascular disease categories, reflecting the clinical contexts most commonly linked to these lipid-associated proteins (**Tables S3**, **S4**).

The cytoskeleton-related theme (actin binding, actin filament organization, supramolecular fiber organization) spans VCL, PFN1, VASP, FLNA, CTTN, and related proteins, pointing to cytoskeletal organization and structural dynamics. Tissue annotations for this module center on blood and blood platelets, with additional representation in vascular-associated contexts and smooth and cardiac muscle, consistent with mechanically responsive and contractile tissue environments. Disease terms linked to this set include thrombosis-related, fibrolytic and cardiovascular disease categories, together with broader disease annotations associated with cytoskeletal organization.

A third cluster is characterized by enrichment for platelet activation and aggregation, acute phase response, and defense response, driven by proteins including FGA/FGB, PF4/PPBP, ITIH4, SAA family members, SERPINA1, and CFD. This module is annotated to blood, blood plasma, blood platelets, and liver tissues, consistent with circulating plasma proteins, platelet biology, and hepatic acute-phase production. Disease associations for this block include systemic inflammatory disease, coagulation-related disease, and cardiovascular disease categories, matching the processes highlighted by the GO annotations.

Overall, **PPA 526** (**Figure 3A**) can be used to follow the main biological themes at a lower level of complexity, while **PPA 1514** (**Figure 3B**) can be used to broaden coverage and sense proteoforms that participate across multiple biological systems. As annotation resources continue to improve, these views can be extended to include proteoform-aware information, allowing coverage to be evaluated not only by the presence of a gene/protein-level response, but by which proteoforms are detected and whether they are likely to be functional in the biological systems of interest.

Analysis of post-translational modifications (PTMs) across proteins stratified by biological processes revealed clear, process-specific PTM signatures. Proteins involved in cytoskeletal organization and actin filament dynamics were predominantly associated with regulatory PTMs, most notably serine/threonine phosphorylation, along with α-amino acetylation and lysine methylation (**Figure S4**). These PTMs are characteristic of intracellular and platelet derived proteins that undergo rapid, reversible regulation during cytoskeletal remodeling and platelet activation. The enrichment of phosphorylation in these pathways is consistent with kinase driven control of actin dynamics and highlights the tight coupling between signaling and structural reorganization in these processes.

In contrast, proteins associated with lipid transport, lipoprotein remodeling, and cholesterol efflux exhibited a markedly different PTM landscape dominated by truncation, deamidation of asparagine and glutamine, methionine oxidation, and N terminal pyroglutamate formation (**Figure S5**). These PTMs are typical of long-lived, secreted plasma proteins such as apolipoproteins and likely reflect cumulative chemical modification, proteolytic processing, and oxidative exposure in circulation rather than active regulatory signaling. Proteins involved in platelet activation and inflammatory responses displayed a mixed PTM profile, combining phosphorylation-based regulatory modifications with truncation and oxidative changes (**Figure S6**), consistent with their dual intracellular origin and extracellular functional roles. Overall, these results demonstrate that PTM distributions align closely with biological function and protein localization, supporting the biological relevance of proteoform-level sampling across PPA panels.

### Exploratory Streamlined Quantitative Evaluation of PPA Panels in Sets 2 and 3

**PPA 526** was applied to a cohort comprising individuals carrying a SERPINE1 mutation (**Set 2**, n = 27; see **Figure 4**). In this analysis, the soluble fraction was cleaned and analyzed by mass spectrometry without FAIMS. **PPA 1514** was applied to a second cohort composed of patients with hepatic cirrhosis (**Set 3**, n=30) across three disease stages: compensated cirrhosis (n=9), compensated with portal hypertension (n=10) and decompensated cirrhosis (n=11), with proteoform analysis performed using FAIMS prior to single ion proteoform detection. We hypothesized that most proteoforms included in the panels could also be quantified in **Sets 2** and **3**, given the sample diversity and analytical depth used to define the panels. As proof of concept for potential applications, we primarily focused on describing proteoform identifications, panel coverage, reproducibility, and the main differences observed among the studied groups. Thus, the results below include an overview of biological processes associated with these changes, presented in a descriptive and exploratory manner, without advancing interpretative claims on disease biology.

**Figure 4.**
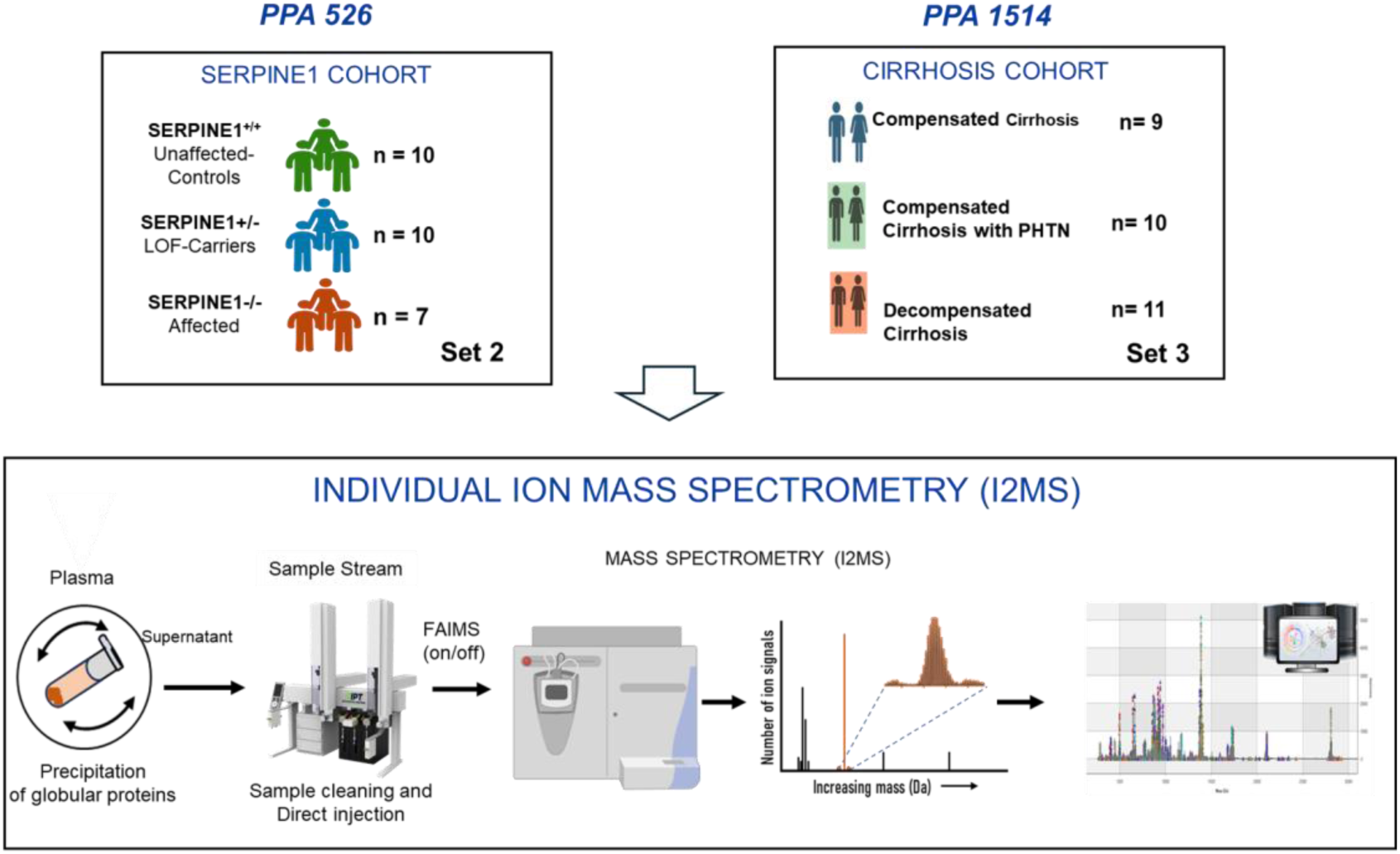
General workflow for Proteoform Plasma Analysis (PPA) using direct injection and Individual ion mass spectrometry (I²MS). We quantified proteoforms from panels **PPA 526** and **PPA 1514** in **Set 2** and **Set 3,** respectively. A soluble fraction after acetonitrile precipitation (SPAP fraction) was applied to deplete high-abundance proteins for top-down mass spectrometry. **Panel 526** was applied to an independent cohort that included individuals carrying a SERPINE1 mutation (**Set 2**), while **PPA 1514** was applied to a cirrhosis cohort (**Set 3**). Analyses of **Sets 2** and **3** followed the same analytical strategy but with FAIMS deactivated (off) for **Set 2** and activated (on) for **Set 3.**

### The SERPINE1 cohort of resilient agers (**Set 2**)

The sample **Set 2** containing 27 subjects were run using the **PPA 526** panel (**Figure 4, top left**). A total of 452 of 526 targeted proteoforms were detected, reflecting expression from 46 of 59 genes targeted in the panel. This corresponds to 86% coverage of the targeted proteoforms and 73% coverage of the corresponding genes/proteins. Despite being derived from a limited discovery set (81 individuals), the database supports robust detection of the majority of targeted proteoforms in an unrelated sample set, demonstrating that its scope is adequate for application to independent cohorts. The proteoform mass distribution for **Set 2** exhibited trends like those observed in **Set 1** (**Figure 5B**), with the majority of detected species below 20 kDa, in agreement with previous results for the acetonitrile-soluble plasma proteome^23^. The number of proteoforms quantified per sample in **PPA 526** on this cohort was ∼430 proteoforms per sample, with a coefficient of variation for the number of unique proteoforms identified of 0.54% (**Figure S7A**).

**Figure 5.**
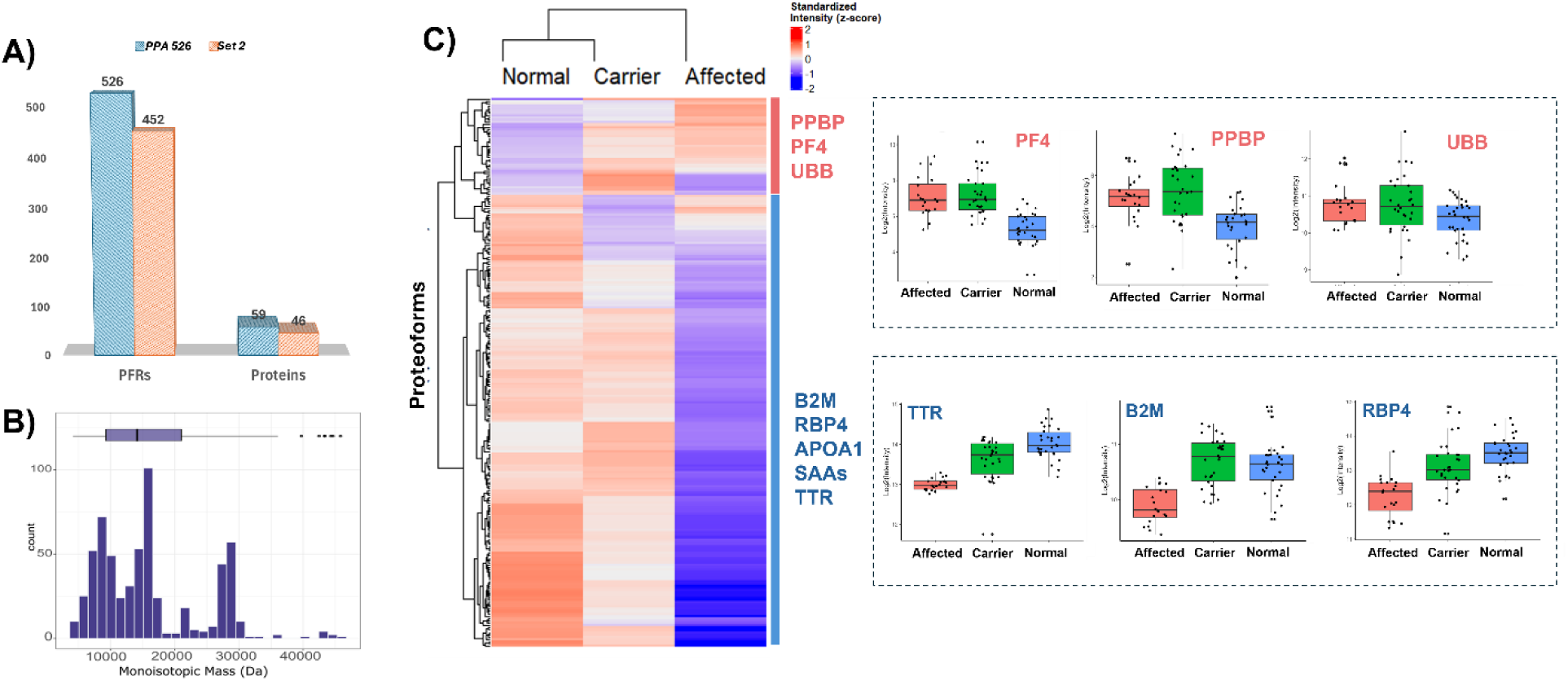
Proteoform quantification in an independent study of a cohort in the context of aging (Set 2). **A)** Bar plot summarizing PFR and protein identifications in **Set 2**, corresponding to **PPA 526**. **B)** Mass distribution of proteoforms identified in **Set 2**. **C)** Heatmap of differential expression analysis (DEP) in **Set 2**, demonstrating distinct proteoform expression patterns across the three experimental groups. Selected box plots highlighting PFRs of interest are shown on the right.

Quantification results revealed notable differences among the three groups in this cohort. Individuals heterozygous for PAI-1 are known to have genetically mediated protection from biological aging (reported here as the PAI-1/SERPINE1 cohort). Major changes were observed in the expression levels of proteoforms in individuals carrying a heterozygous mutation in the SERPINE1 gene (Affected) compared with the other groups (**Figure 5C** and **Figure S7B**). Two distinct patterns emerged: one set of proteoforms was downregulated in the Affected group, while another set was upregulated. For example, proteoform families encoded by PPBP, PF4, and UBB mainly showed elevated expression in affected individuals, with a more pronounced increase compared to the Normal group (**Figure 5C**). Conversely, proteoforms from TTR, APOA1, RBP4, B2M, CST3 and SAAs proteins were mainly downregulated in the Affected group, displaying trends similar to those when compared with the Carrier or the Normal group (**Table S5**).

These differences suggest a baseline of lower chronic inflammation in individuals carrying a heterozygous mutation in the SERPINE1 gene. This is supported by reduced levels of SAA1-2 and, more notably, a decrease in SAA4 proteoforms, a constitutive protein associated with chronic inflammation rather than acute-phase responses ^50^. Similarly, lower expression of B2M may indicate reduced immune activation and lymphocyte turnover, further supporting a state of low chronic inflammation those with a heterozygous SERPINE1 gene ^51, 52^. Given that PAI-1 promotes inflammation by enhancing immune cell infiltration and activating pro-inflammatory cytokines ^53^, a mutation in SERPINE1 leading to PAI-1 deficiency would be expected to attenuate chronic inflammatory signals in individuals not experiencing infections or acute-phase reactions.

In addition, signals of homeostatic responses were observed in the Affected group, including changes associated with HDL regulation (increased levels of certain APOA1 proteoforms) and platelet activation (elevated PF4 and PPBP). Interestingly, some proteoform families showed downregulation specifically in the Affected group, without significant changes between the Normal and Carrier groups (**Table S5**). These included proteoforms from families encoded by B2M, SAA4, APOA1, and certain APOA2 proteoforms, particularly those dominated by the dimer form truncated on the C-terminus (**Figure S8**). These proteoforms could be of particular interest, considering the physiological differences observed in the Carrier population compared to the Affected group. Notably, the Carrier group represents a population characterized by unique demographic and lifestyle factors, including extended longevity and reduced incidence of certain age-related diseases ^54^. This aging-related context suggests that the observed proteoform patterns, especially those showing divergence between Carrier and Affected individuals, may reflect adaptive mechanisms or protective molecular signatures associated with healthy aging and potentially sense patterns consistent with inflammaging ^55^. Interestingly, APOA2 proteoforms, like APOA1, displayed heterogeneous behaviors, suggesting possible functional proteoform diversity (**Figure S8**).

Extending the results observed above, a cohort of cirrhosis subjects organized in sample **Set 3** used **PPA 1514** to detect 1354 of the 1514 targeted species across the 118 of 155 total genes encoding these proteoforms. This corresponds to 89% proteoform coverage and 76% gene/protein coverage. These results demonstrate substantial recovery of the targeted proteoform space and support the use of reference proteoforms derived from this study for quantitative analyses across diverse human plasma samples.

The PFR mass distribution followed the trend of the panel (**Figure 6B**), and proteoform quantification exceeded 1,300 proteoforms per technical replicate (**Figure S9A**). **This level of resolution enabled clear separation of cirrhosis stages by principal component analysis (PCA)** (**Figure 6C**). These groups comprised samples from patients representing three progressive stages of liver disease: compensated cirrhosis (with and without portal hypertension) and decompensated cirrhosis (**Figure S9B**). We previously demonstrated that using LC–MS-based proteoform analysis, proteoform dynamics across these patient populations enable partial stratification of cirrhosis stages based on distinct proteoform signatures ^23, 48^. Here, by quantifying PFRs from the **PPA 1514** panel in a small cohort of 15 cirrhosis patients, we demonstrate that these proteoform dynamics are reproducible and sensitive to underlying biological differences. Notably, similar molecular functions previously associated with disease stage–specific differences were again identified^23, 48^ (**Figure 6D, Table S6**). These functions were primarily related to the plasma lipoprotein particle system, driven by proteoforms originating from APOA1, APOA2, and APOC3, with additional contributions from abundant plasma proteins involved in transport and coagulation, including haptoglobin, fibrinogen alpha chain, and transthyretin. Peptidase activity–related processes were likewise enriched and were represented by proteoforms from cysteine and serine protease inhibitor families, such as cystatin C and SERPINA1.

**Figure 6.**
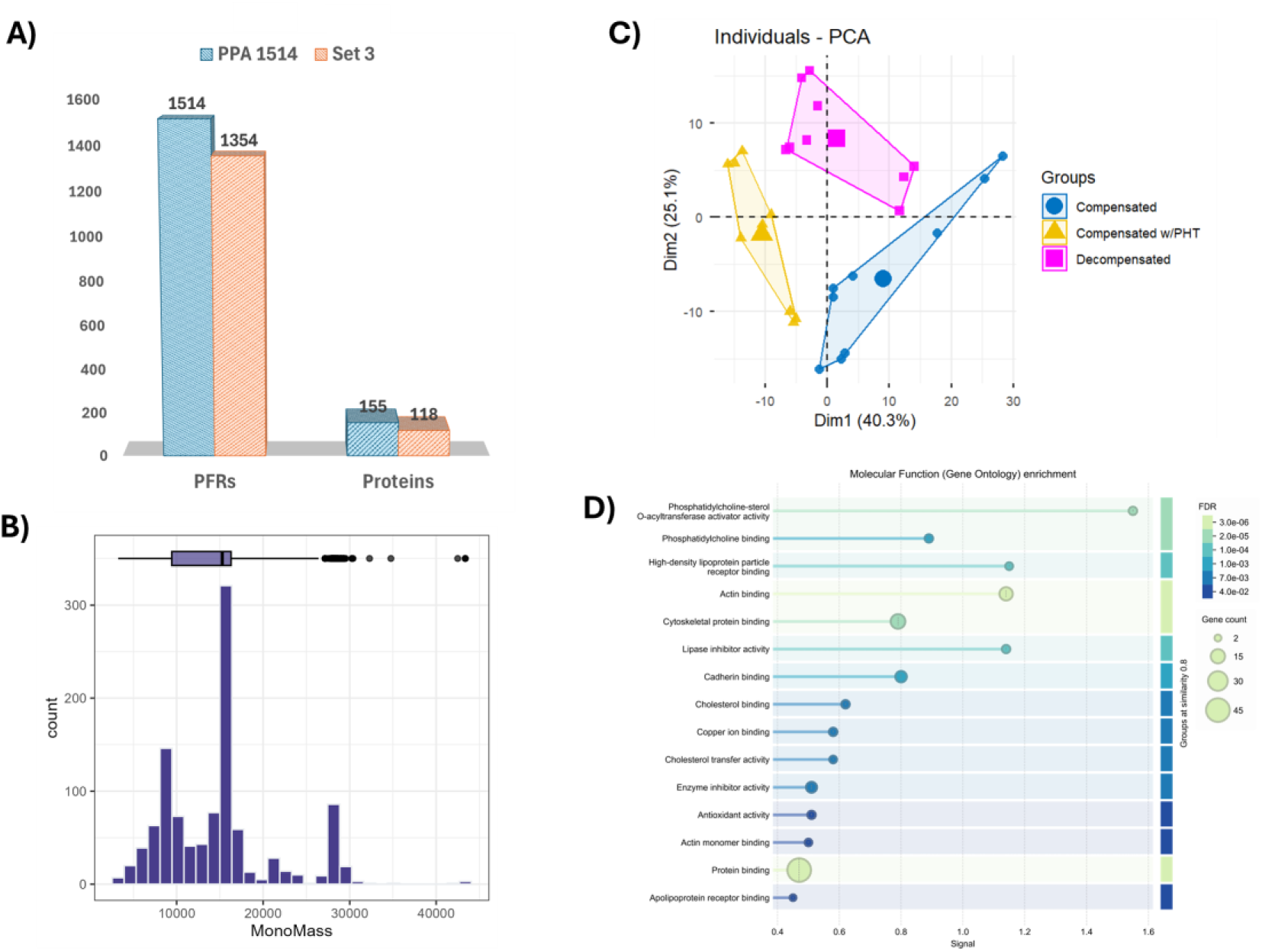
Proteoform quantification in an independent cirrhosis cohort (Set 3). (**A)** Bar plot summarizing PFR and protein identifications in Set 3, corresponding to **PPA 1514**. **(B)** Mass distribution of proteoforms identified in Set 3. **(C)** Principal component analysis (PCA) using components 1 (X-axis) and 2 (Y-axis), demonstrating distinct proteoform expression patterns across the three experimental groups. **(D)** Top 10 molecular functions enriched by DEP from the current cirrhosis study.

### Reproducibility of quantification is based on single ions

We previously demonstrated that the coefficient of variation (CV) for the acetonitrile precipitation process, combined with proteoform label-free quantification from LC-MS data, was consistently below 20% ^23^. Here, we hypothesized that single-ion quantification following direct plasma proteoform analysis by I²MS would exhibit comparable reproducibility.

The median technical variation observed across the three groups in **Set 2** following application of **PPA 526** remained consistently below 20% of CV (**Figure S10A**). More than 58% and 69% of PFRs exhibited CVs below 20% and 25%, respectively, demonstrating good reproducibility across technical replicates (**Table S7**).

Similarly, analysis of **Set 3** using **PPA 1514** yielded technical CVs consistently below 15% with more than 67% and 76% PFRs with CVs below 20% and 25%, respectively, reflecting good analytical reproducibility following incorporation of FAIMS (**Figure S10B, Table S8**). For the reference sample, the median CV was below 10%, with over 80% and 90% of PFRs displaying CVs below 20% and 25%, respectively (**Figure S10C, Table S9**). Collectively, these results demonstrate that single-ion measurements enable reproducible quantification of targeted plasma proteoforms across both panels and are broadly consistent with the performance reported for highly accurate peptide MS-based multiplexed proteomics workflows used for plasma protein quantification^11^.

In large scale, label free proteomics studies, median coefficients of variation (CVs) below approximately 20–25% are commonly considered indicative of good or acceptable quantitative reproducibility at the peptide level, particularly when analyzing complex biological matrices and extended cohorts, where additional sources of biological and technical variability are expected ^11, 15, 56–58^. At the proteoform level, quantitative reproducibility can be more challenging than peptide-based measurements, as multiple molecular forms arising from sequence variation, post-translational modification, and proteolytic processing contribute to increased signal complexity and variability, particularly in complex biofluids, necessitating context dependent interpretation of CVs ^57, 59, 60^.

### Comprehensive analysis of PPA 1514 and PPA 526

To contextualize the plasma proteome coverage of the **PPA 1514** and **PPA 526** panels, we first evaluated their protein abundance profiles using concentration estimates reported in the Human Protein Atlas (HPA), see **Table S11**. When ranked by abundance, both PPA panels show a clear enrichment toward the more abundant fraction of the plasma proteome, with a substantial proportion of panel proteins falling within the Top 100 and Top 1000 most abundant plasma proteins (**Figure S11, Table S11**). Compared with the Olink 3K panel ^40^ (**Tables S11** and **S12**), which spans a broader dynamic range, PPA panels preferentially target higher abundance proteins, consistent with their design and analytical sensitivity. This observation highlights the complementary nature of affinity-based panel selection strategies and their impact on proteome coverage.

A quantitative comparison of abundance rankings reveals that **PPA 526** consistently captures a higher fraction of proteins within the Top 100 category than **PPA 1514**, whereas both panels converge more closely at the Top 1000 level. This pattern suggests that **PPA 526** may be particularly optimized for detecting proteins at the upper end of the plasma concentration spectrum, while **PPA 1514** extends further into lower abundance regions. In contrast, proteins from the Olink 3K panel populate the Top 100 category to a much lesser extent, reflecting the deliberate inclusion of low abundance signaling and regulatory proteins within that platform. Together, these findings illustrate how panel composition and assay sensitivity influence detectable abundance ranges (**Figure S11**).

The absolute abundance of classification further reinforced these trends. When proteins were grouped into high (>40 µg/mL), medium (10 ng/mL–40 µg/mL), and low (<10 ng/mL)^15, 17, 23,61^ abundance categories based on estimates from HPA BUP mass spectrometry, the majority of HPA reported proteins—and particularly those represented in the Olink 3K panel—fell into the low abundance category (**Table S11-S12**). In contrast, **PPA 1514** and **PPA 526** displayed a relatively higher proportion of medium and high abundance proteins compared to the representation in the lowest concentration range (**Table S11**). Importantly, proteins shared between Olink and PPA panels were enriched in the medium abundance group, suggesting that overlap between platforms is most prominent for proteins present at intermediate plasma concentrations (**Figure S12**).

Functional categorization of shared proteins provided further insight into panel complementarity. Among the 70 proteins common to **PPA 1514** (**Table S13**) and Olink 3K, functional annotations derived from Olink taxonomy revealed a predominance of inflammation and cardiometabolic related proteins, followed by neurological and oncological categories (**Figure S13**). This functional skew consists of the biological roles of moderately to highly abundant plasma proteins, which often participate in systemic signaling, immune response, and homeostasis. Expansion of this analysis to proteoform representations demonstrated a marked amplification of inflammatory associated proteoforms, underscoring the added biological resolution captured when proteoform diversity is considered alongside basic protein identity.

Taken together, these analyses demonstrate that **PPA 1514** and **PPA 526** provide robust coverage of medium to high abundance plasma proteins while retaining meaningful overlap with the more expansive and low abundance focused Olink 3K panel. Importantly, shared proteins between PPA and Olink panels span functionally relevant pathways and abundance ranges that are well positioned to assess the difference between proteins and proteoforms in joint clinical studies.

### Acquisition-Time–Driven Throughput and Proteoform Coverage in FAIMS-Enabled Plasma Profiling

To assess the effect of sampling time, we ran the **PPA 1514** panel at increasingly lower data acquisition times. For each of the 1, 5, 10, 20, and 40 minutes of time allocated, the key result is the number of PFRs that could be quantified. Importantly, the **PPA 1514** panel not only increased proteoform and protein coverage relative to **PPA 526** but also enabled substantial reductions in acquisition time, from approximately 50 minutes to as little as 20, 10, 5, or even 1 minute, while maintaining comparable identification performance. To systematically evaluate FAIMS-enabled quantification across these regimes, proteoform ions were analyzed from a reference plasma sample using acquisition times ranging from 1 to 40 minutes. Even with a 1 minute acquisition, 597 proteoforms could be quantified per run, with coverage increasing to 1072, 1100, 1250, and 1272 proteoforms per sample at 5, 10, 20, and 40 minutes, respectively (**Figure S14**).

As data acquisition times for **PPA 1514** dropped, the median CVs for PFR quantitation remained consistently below 20% across all acquisition times, underscoring the potential of PPA for robust and scalable reproducibility (**Table S10** and **Figure S10D**). As shown in the density plot of R² values (ion count versus acquisition time), the vast majority of proteoforms exhibited near-perfect linear behavior. The experiment demonstrated excellent linearity in proteoform quantification as a function of acquisition time with a median R² of 0.995 for proteoforms quantified across time points (**Figure S10E**).

The results for dropping data acquisition times demonstrate that high proteoform identification can be retained across a broad range of acquisition times (**Figure S14).** To contextualize the throughput implied by these acquisition times, samples per day (SPD) performance was estimated and compared with recent BUP workflows for plasma proteomics, reporting approximately 100 SPD using ∼11.5 minutes gradients ^13^. Assuming continuous instrument operation, an 11.5-minutes acquisition time corresponds to roughly 125 samples per day, a value commonly operationalized as 100 SPD after accounting for equilibration and instrumental overhead. Within this framework, the identification depth achieved by **PPA 1514** supports throughput levels comparable to, or exceeding, those reported in highly engineered BUP workflows using LC-MS. Specifically, 20 minutes acquisitions correspond to ∼72 SPD while yielding ∼1250 proteoforms per sample, placing this regime below the nominal 100 SPD threshold but with substantially greater proteoform depth. At 10 minutes PPA times, projected throughput increases to ∼144 SPD while preserving >1100 quantified proteoforms per run, exceeding the effective throughput of 11.5 minutes workflows. Shorter acquisitions further extend this trend, with 5 minutes. PPA run times corresponding to ∼288 SPD and still yielding >1000 proteoforms, and 1 minute acquisitions enabling a projected throughput of >1400 SPD, with nearly 600 quantified proteoforms per sample (**Figure S14**).

Notably, these throughput estimates are conservative, as **PPA 1514** targets up to 1514 proteoforms and the analysis was performed using a single reference plasma sample, suggesting that additional proteoform coverage may emerge across more diverse sample cohorts. Collectively, these results indicate that **PPA 1514** enabled by FAIMS supports proteoform identification depth compatible with 100 SPD class workflows at acquisition times of 10 minutes or less, partially mapping the quantitative tradeoff between acquisition time and analytical depth.

In addition to the deeper coverage of **PPA 1514** workflow, we evaluated an even more streamlined strategy, termed **PPA 232**, designed to maximize throughput while minimizing sample preparation complexity. **PPA 232** employs the same FAIMS-enabled acquisition strategy as **PPA 1514** but omits ACN precipitation, relying instead on simple plasma dilution followed by direct mass spectrometric analysis. Under these conditions, **PPA 232** enabled quantification of 54 proteins corresponding to **232** PFRs in a single shot measurement (**Table S14**). Despite reduced analytical depth relative to **PPA 1514, PPA 232** captured a conserved plasma proteoform signature (**Figure S15 A-B**), exhibiting >90% overlap at the PFR level and ∼99% overlap at the protein level with the larger panel (**Figure S16A-C**). Quantitative reproducibility under these high throughput conditions was robust. Across replicate measurements, **PPA 232** achieved a median coefficient of variation of ∼14%, with approximately 75% of PFRs exhibiting CVs <20% and ∼90% below 25%, demonstrating consistent proteoform level quantification across short runs and extended acquisition series (**Figure S16D**).

While ACN precipitation in **PPA 1514** substantially enhances proteoform discovery and functional resolution, MS high-throughput operation is fundamentally an acquisition time problem, not a sample preparation constraint. This establishes FAIMS-enabled plasma proteoform analysis as a continuum spanning ultra-high throughput, single shot signature measurement to deep, discovery-oriented profiling, depending on experimental objectives. Collectively, the data reported here indicate that I^2^MS- and FAIMS-enabled acquisition supports proteoform identification depth compatible with 100 SPD class workflows at acquisition times of 10 minutes or less. While sustained high throughput operation was not directly assessed in this study, the observed tradeoff between acquisition time and proteoform coverage defines a quantitative framework suggesting the feasibility of high throughput, proteoform resolved plasma profiling under appropriate experimental conditions.

### Limitations

Although we report proteoform results from just over 100 individuals, this sample size remains limited and should be expanded to demonstrate generalizability. While we developed a pipeline for MS1 feature detection and quantification using a curated MS2 spectral database from similar samples, the current results depend on accurate MS1 feature detection without formal FDR control. Functional enrichments and annotations were performed at the protein (gene) level, whereas the experimental measurements were obtained at the proteoform level. Therefore, differences among individual proteoforms, such as truncations, post-translational modifications, or processing variants, are not explicitly captured in the enrichment analyses. This abstraction can underestimate or obscure the biological relevance of specific proteoforms, particularly for proteins such as hemoglobins, whose proteoform diversity is frequently observed in clinical serum or plasma samples^23, 24, 48^. Proteoform specific signals may reflect a mixture of biological variation and pre-analytical factors, including sample handling, plasma processing, or hemolysis, which can confound downstream interpretation if not carefully considered.

In this study, hemoglobin proteoforms were included in the PPA panels to enable tracking of these species and to reduce the risk of false positive identifications or mismatches across samples. However, because the presence of hemoglobin proteoforms in plasma may predominantly stem from sample handling related artifacts rather than disease specific biology, they were excluded from downstream enrichment and interpretative analyses. While proteoform resolved information can provide meaningful biological insight in contexts where clinical metadata or study design support such interpretations, the present analyses intentionally adopt a conservative strategy to avoid overinterpretation of signals likely driven by pre-analytical variability. Some proteoforms may be partially or completely nonfunctional compared to their intact counterparts, and this functional heterogeneity is not reflected in current annotation databases. Future work will aim to bridge this gap by evaluating how proteoform specific measurements contribute to biological system coverage, enabling a progression from simpler, gene centric models toward more complex and realistic representations of biological function. As our main objective is to apply this proteoform panel to large cohorts for detecting substantial proteoform changes, we plan to validate key findings through subsequent fragmentation (LC-MS/MS and I²MS²)^31, 62^ or other orthogonal and complementary techniques.

## Conclusion and Outlook

The concept of PPA can be adapted to other fluids (CSF, serum) and plasma from other organisms, yet the top challenges are also scalability and comparing statistical power and effect sizes relative to protein-only plasma assays. Olink-based plasma proteomics has scaled dramatically, moving from studies of thousands of individuals with thousands of proteins, reflected in the UK Biobank effort, profiling over 50,000 participants for ∼3,000 proteins ^7^. This progression marks a major step toward increased coverage of the plasma proteome and can align with proteoform-aware PPA platform, where population-scale studies are now feasible. As proteoforms are more proximal to phenotypes, they should provide a larger effect size ^3, 22^, and require smaller cohorts to sense human biology across a population and the state of an individual ^22^. With a larger effect of size from detecting proteoforms, combined with expanding proteoform coverage (*e.g.*, through use of Extracellular Vesicles or improved I^2^MS), deeper analyses are clearly possible in industrial, academic, and regulatory settings.

## Data availability

Raw data files are available in the MassIVE repository under accession number MSV000101890. Access credentials will be provided upon request.

## Acknowledgement

The research and optimization of the assay panels reported in this publication were supported by the National Institutes of Health (NIGMS) under Award Number RM1 GM156535. The content is solely the responsibility of the authors and does not necessarily represent the official views of the National Institutes of Health.

## Supporting Information

Supplementary Figures: Provided as a separate file.

Supplementary Tables: https://figshare.com/s/4b92693922ad4f51946c

## Author Contributions

AS contributed to conceptualization, experimental design, sample processing, data collection and data analysis. IP and MH contributed to conceptualization, experimental design and data analysis. KP contributed to experimental design, sample processing, data collection and data analysis. TS contributed to sample processing and data analysis. VW, TDF, ML and NH contributed to experimental design, sample processing, and data collection. CH, PC, DA. PB, TEB, DPL, JTW, DWV contributed to sample collection. EF contributed to conceptualization and experimental design. MAC contributed to conceptualization, experimental design and data analysis. JPM contributed to conceptualization, experimental design, sample processing, data collection and data analysis. NLK contributed to conceptualization, experimental design, initiating lines of technology development, procuring and administering funding, and manuscript revisions. All authors have given approval to the final version of the manuscript. All authors have given approval to the final version of the manuscript.

## Abbreviations

ACN: Acetonitrile
PFR: proteoform
DEP: differentially expressed proteoform
HDL: high-density lipoprotein
LC-MS: liquid chromatography-mass spectrometry
TDP: top-down proteomics
BUP: bottom-up proteomics
HPA: Human Protein Atlas
I^2^MS: Individual Ion Mass Spectrometry
I^2^MS^2^: Individual Ion Mass Spectrometry
SS: SampleStream platform

