## Supplementary Figures for "Targeted Analysis of >1500 Plasma Proteoforms via Individual Ion Mass Spectrometry"

Running Title: Plasma Proteoform Assay (PPA) for >1500 human proteoforms

Keywords: Proteoforms, Plasma, Top-Down Proteomics, Mass Spectrometry, Acetonitrile Precipitation, I2MS, Direct Mass Technology, Charge Detection.

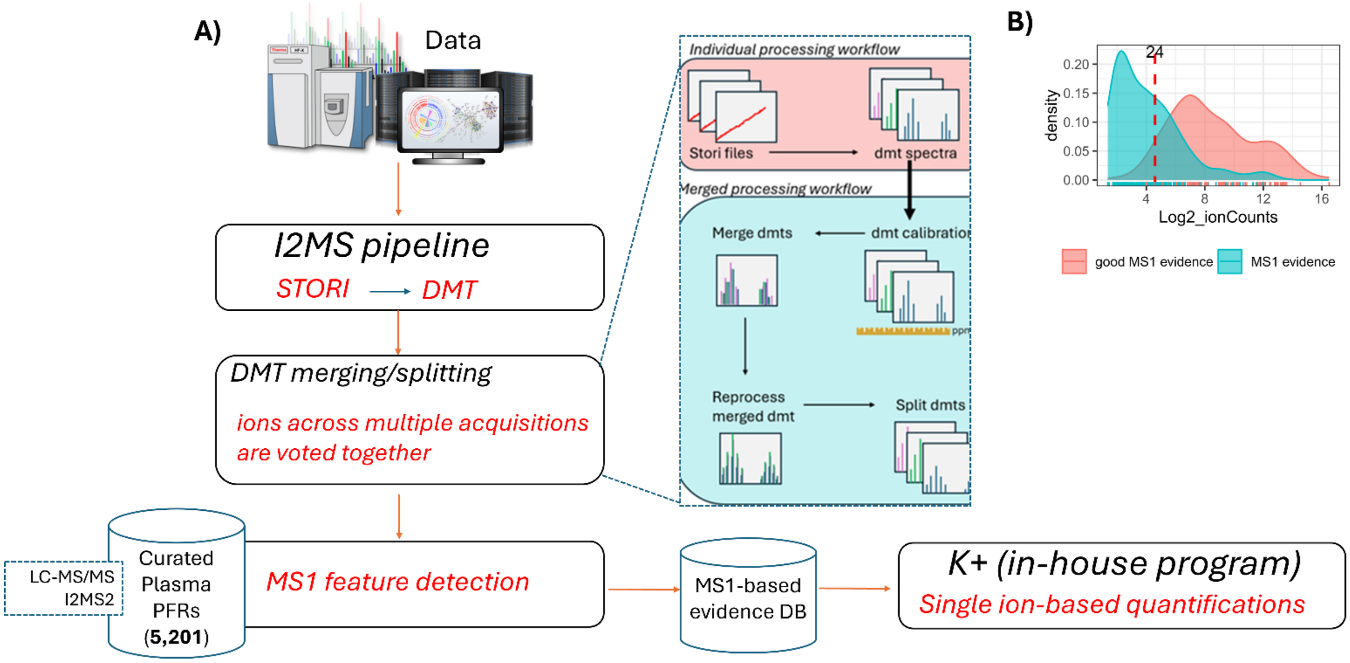

**Figure S1. Extended bioinformatic pipeline used for the identification and quantification of proteoforms by using I2MS analysis. A)** The individual processing workflow consists of the standard processing of 1:1 STORI files to dmt spectra that then undergo MS1 feature detection and assignment and a database of ~5200 proteoforms (PFRs, described in Sanchez *et al.*, 2026). Merged processing involves the added steps of .dmt calibration, merging, reprocessing, and splitting, which allows for ions across multiple acquisitions to be voted together. Individual ions of PFRs are quantified using an in-house program called "K+" . **B)** Distribution of $log_2$-transformed ion counts for proteoforms quantified with (red) and without (teal) good MS1 evidence. Densities illustrate the relative contribution of ions supporting proteoform quantification across datasets. The vertical dashed line indicates the cohort-specific ion count threshold defined as the lower bound retaining 95% of ions associated with MS1-supported proteoforms, which was used to filter proteoforms for downstream analyses.

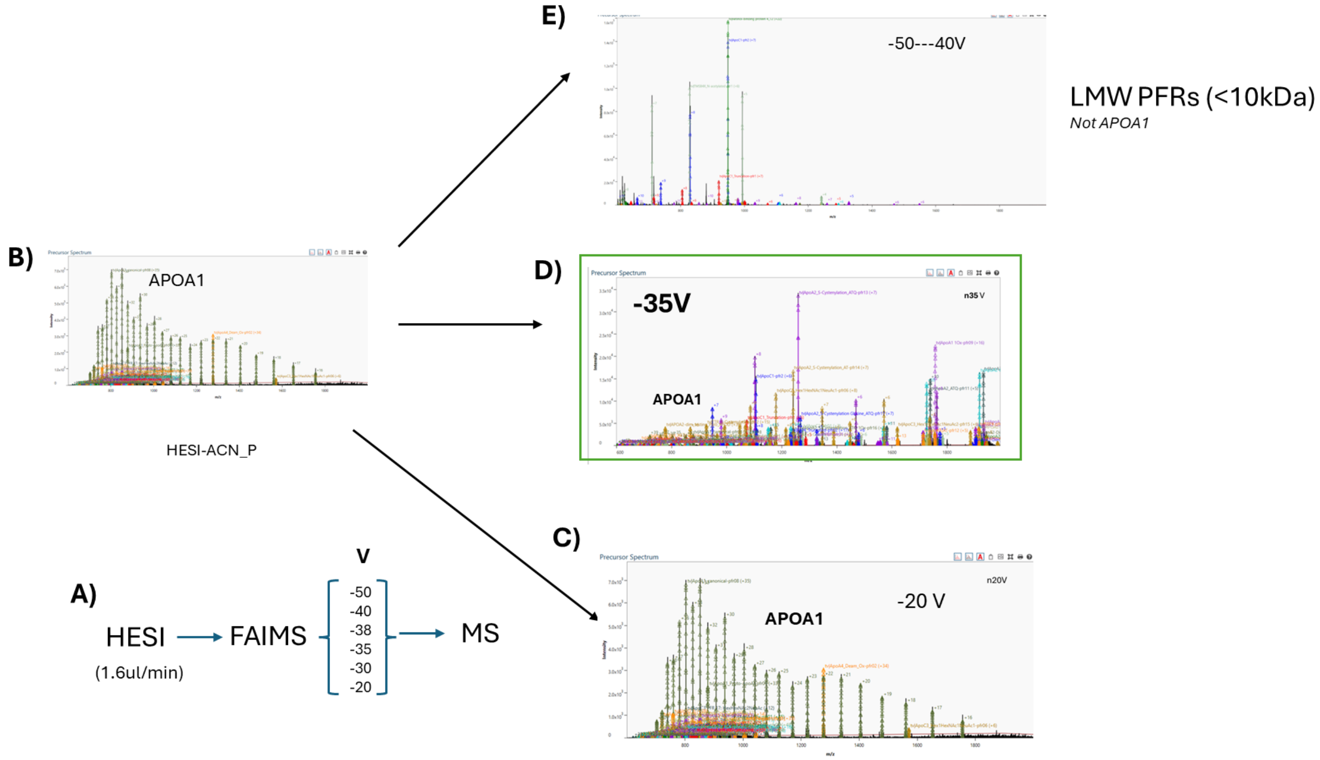

**Figure S2. Optimization of FAIMS compensation voltages for PPA 1514. A)** Overview of the experimental workflow. **B)** Mass spectrum acquired without FAIMS, showing proteoforms from APOA1 as the most abundant proteoform family. **C)** Application of a −20 V compensation voltage (CV) results in APOA1 proteoforms remaining the dominant signals. **D)** Setting the CV to −35 V selectively depletes APOA1 proteoforms, enabling enhanced detection of proteoforms in the 10–30 kDa mass range. **E)** Application of CVs ranging from −40 to −50 V facilitates the identification of low molecular weight (LMW) proteoforms (<10 kDa).

**A)**

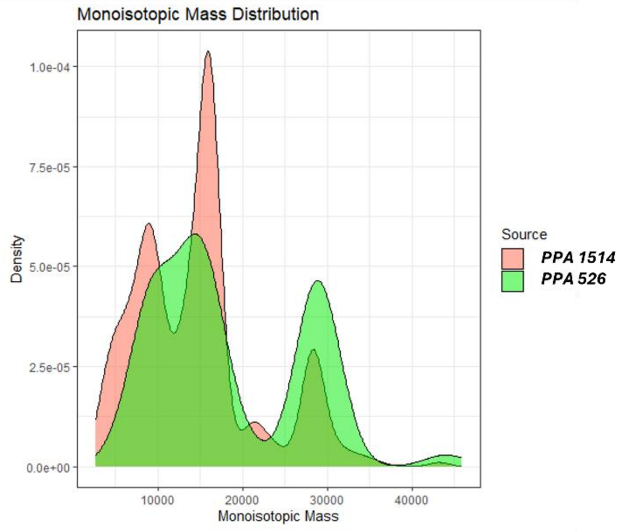

**B)**

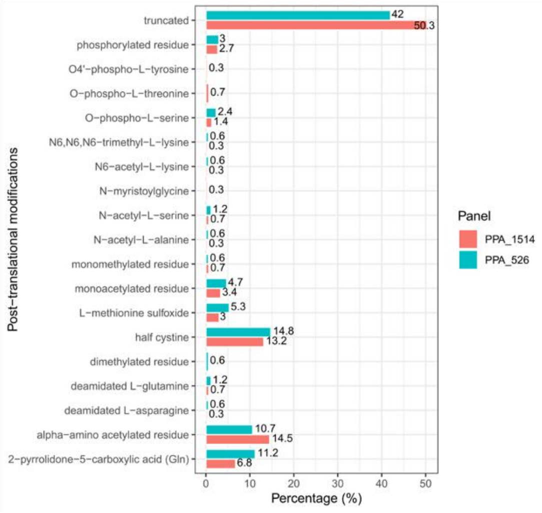

**Figure S3. Distribution of proteoform mass values and PTMs identifications in the two PPA panels. A)** Histograms of proteoform mass distributions, comparing identifications from both panels. **B)** Distributions of PTMs in each PPA panel.

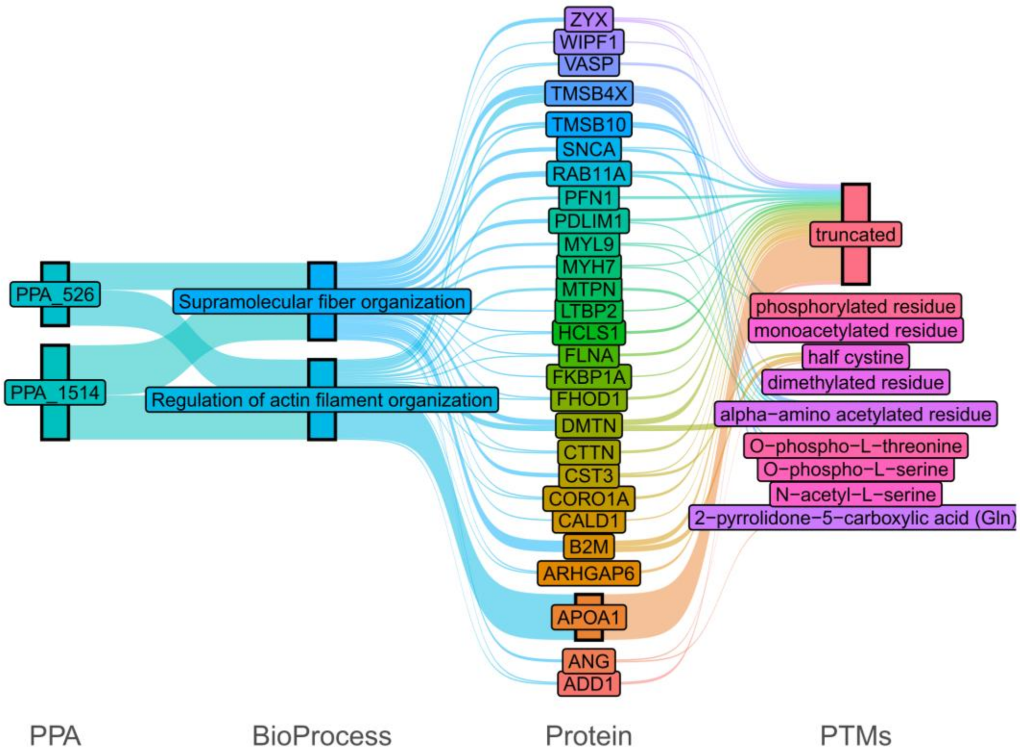

**Figure S4. Distribution of PTMs across categories of biological function.** Panels depict the association between post-translational modifications (PTMs), proteins, and selected cytoskeletal organization and actin filament–related processes, highlighting the proteins involved and their identified PTMs.

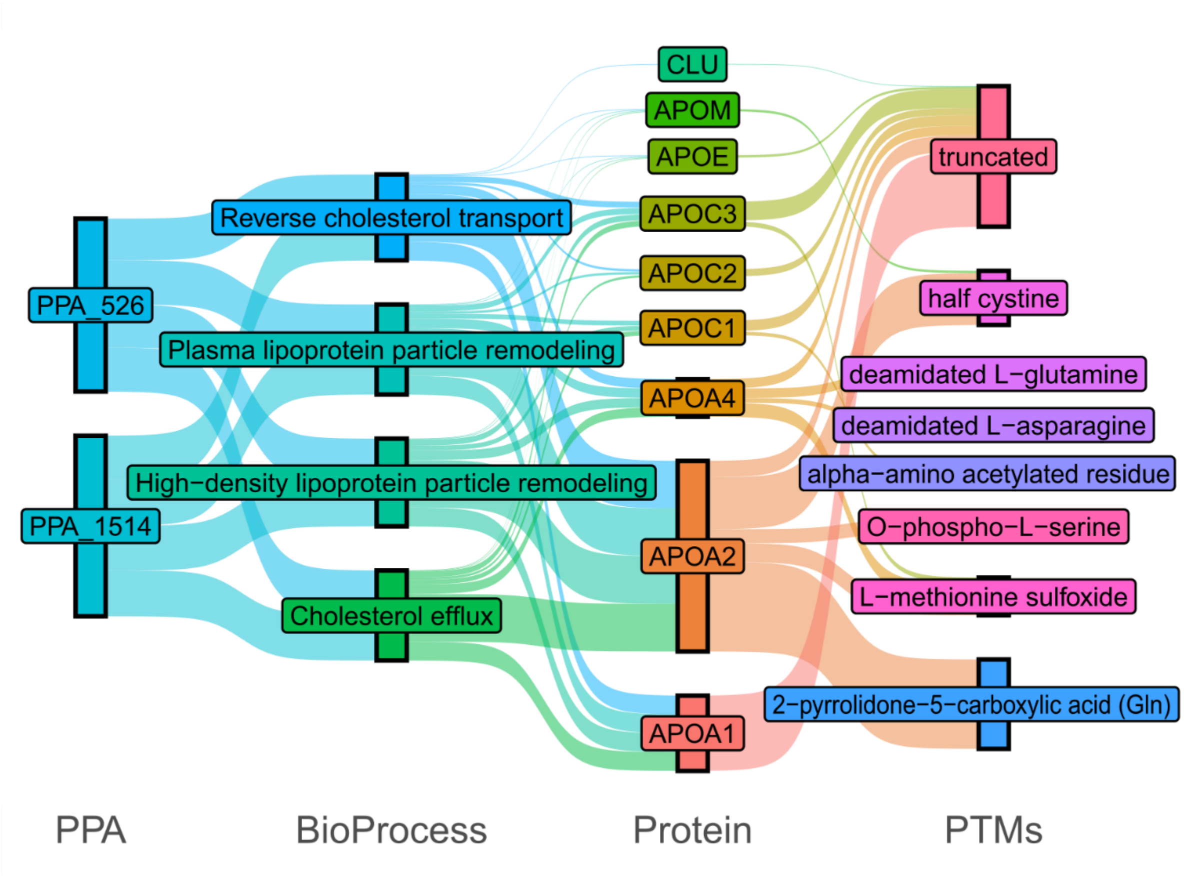

**Figure S5. Distribution of PTMs across categories of biological function.** Panels depict the association between post-translational modifications (PTMs), proteins, and selected lipid transport–related processes and the corresponding proteins associated with PTMs.

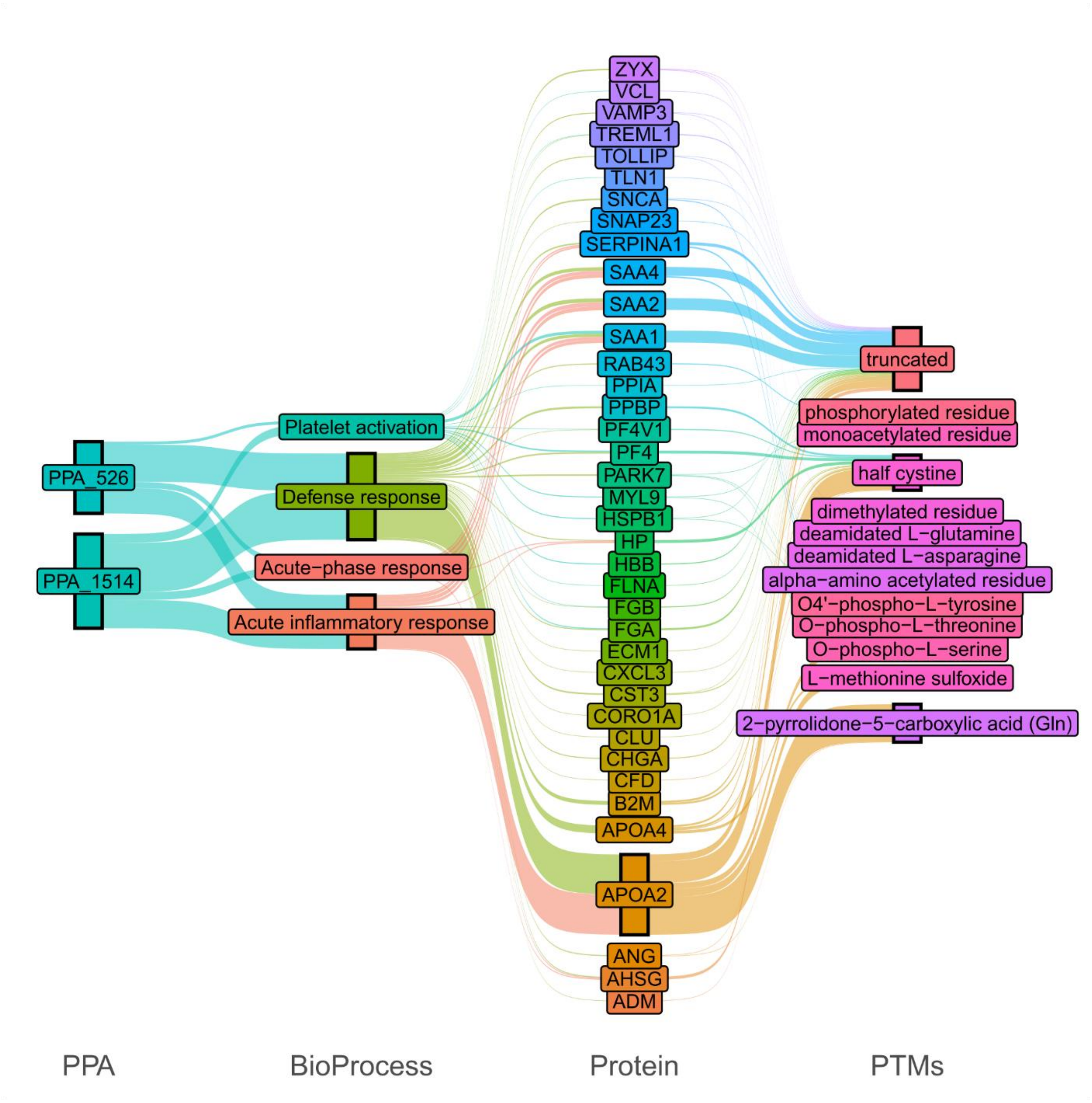

**Figure S6. Distribution of PTMs across categories of biological function.** Panels depict the association between post-translational modifications (PTMs), proteins, and selected platelet activation and inflammatory processes, showing linked proteins and their detected PTMs.

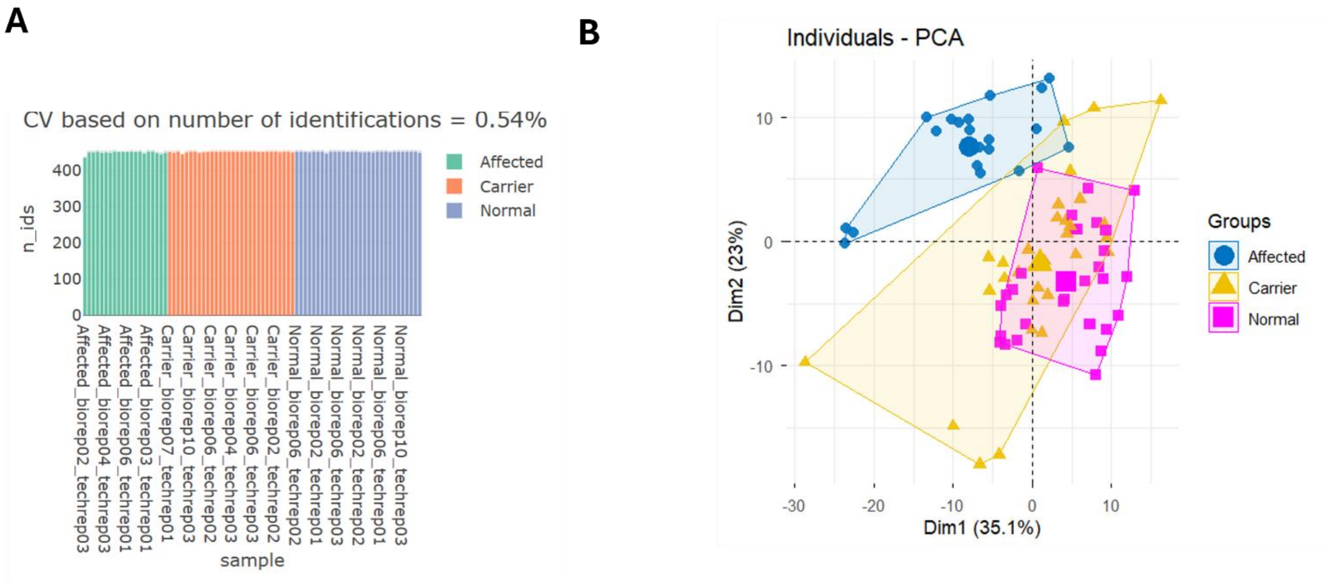

**Figure S7. Quantification of PPA 526 PFRs in the SERPINE1 cohort (Set 2).** Individuals with genetically mediated protection from biological aging (also known as the PAI-1/SERPINE1 cohort). **A)** Number of PFRs identified per sample and corresponding coefficients of variation (CVs) based on identification counts. **B)** Principal component analysis (PCA) using components 1 (X-axis) and 2 (Y-axis).

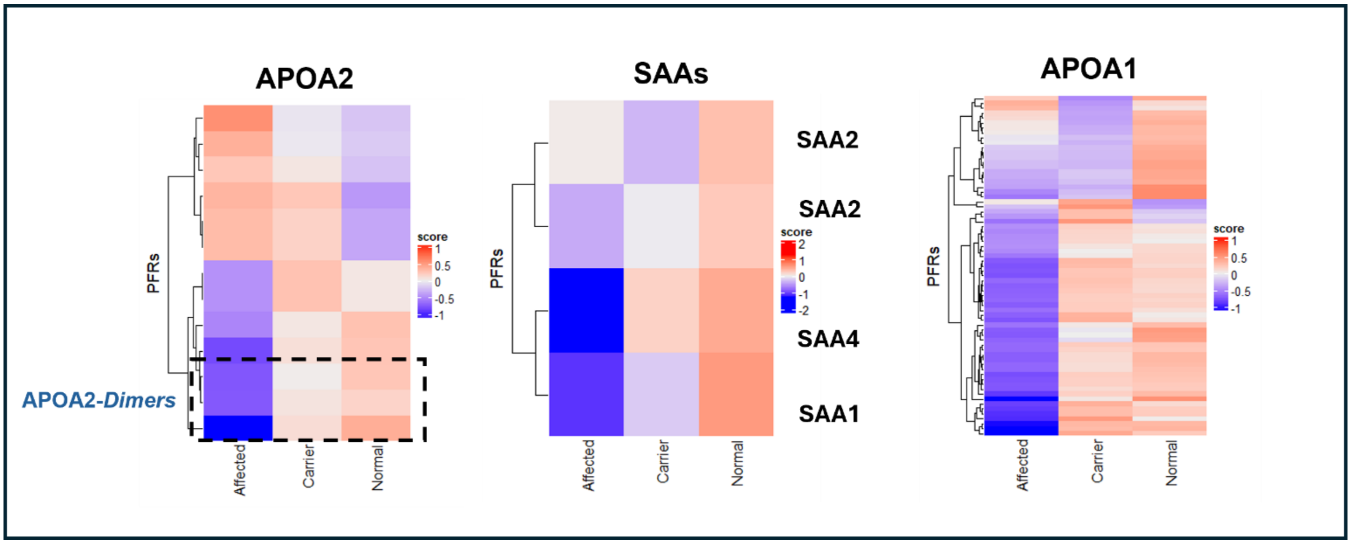

**Figure S8. Dynamics of PFRs from sample Set 2 consisting of subjects with 3 PAI-1 genotypes.** Proteoform responses in Set 2 are predominantly downregulated in the Affected group, while remaining largely unchanged between the Control and Carrier groups. **Left**: APOA2 dimers are downregulated in the Affected group, whereas other APOA2 proteoforms are upregulated. Middle: SAA proteoforms exhibit differential regulation, with SAA4 showing the most pronounced change in the Affected group. **Right**: APOA1 displays heterogeneous behavior overall but is predominantly downregulated in the Affected group.

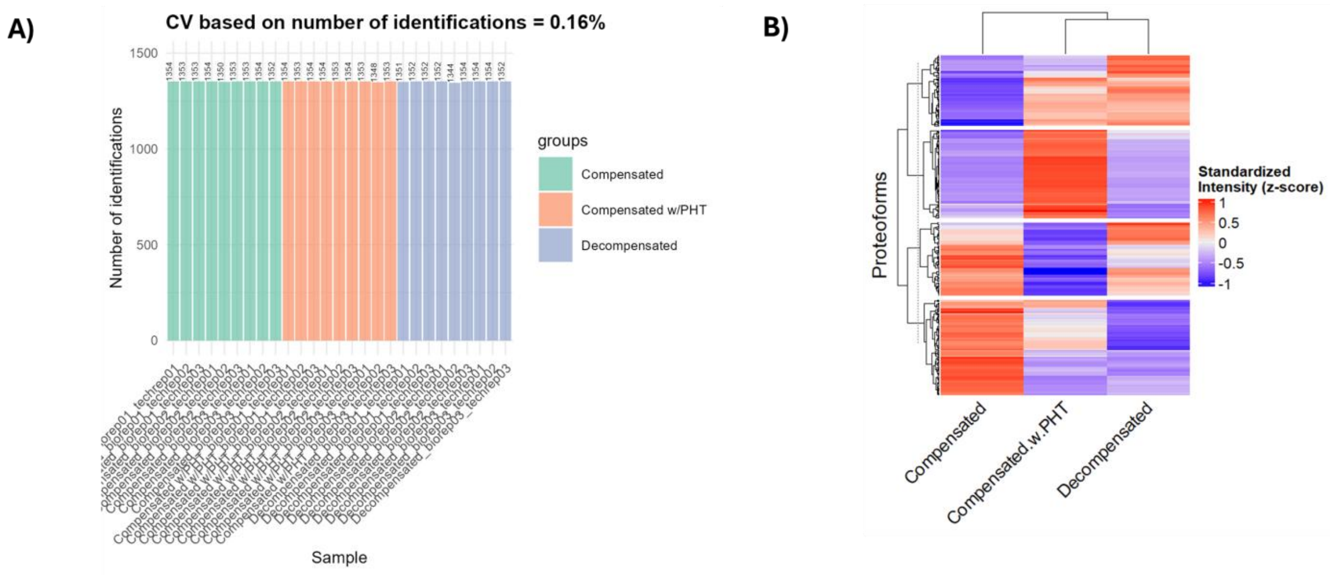

**Figure S9. Quantification of Panel 2 PFRs in the Cirrhosis cohort (Set-3). A)** Number of PFRs identified per sample and corresponding coefficients of variation (CVs) based on identification counts. **B)** Heatmap created from differential expression of proteoform (DEP) analysis in sample **Set 3** from subjects with three different stages of liver cirrhossis, demonstrating distinct proteoform expression patterns across the three experimental groups.

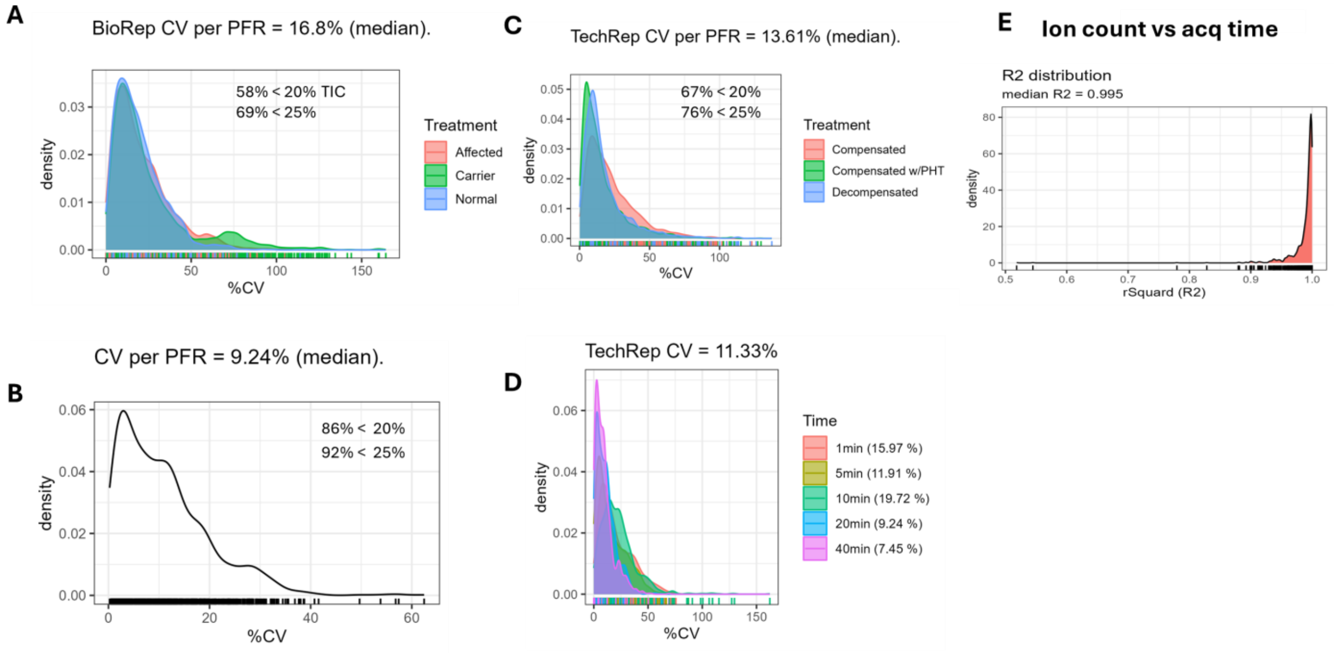

**Figure S10. General reproducibility of direct proteoform analysis using I2MS. A)** Technical coefficient of variation across conditions in Set 2 (**PPA 526**). **B)** Proteoform CVs of ion counts calculated after four technical $I^2$MS injections using a reference sample (**PPA 1514**). **C)** Technical coefficient of variation across conditions in Set 3 (**PPA 1514**). **D)** CVs obtained after detection of proteoforms with different acquisition times (**PPA 1514-** reference sample. **E)** Density plot of distributions of $R^2$ values for PFRs quantified across different acquisition times (1,5,10,20, & 40 min.) from **PPA 1514**.

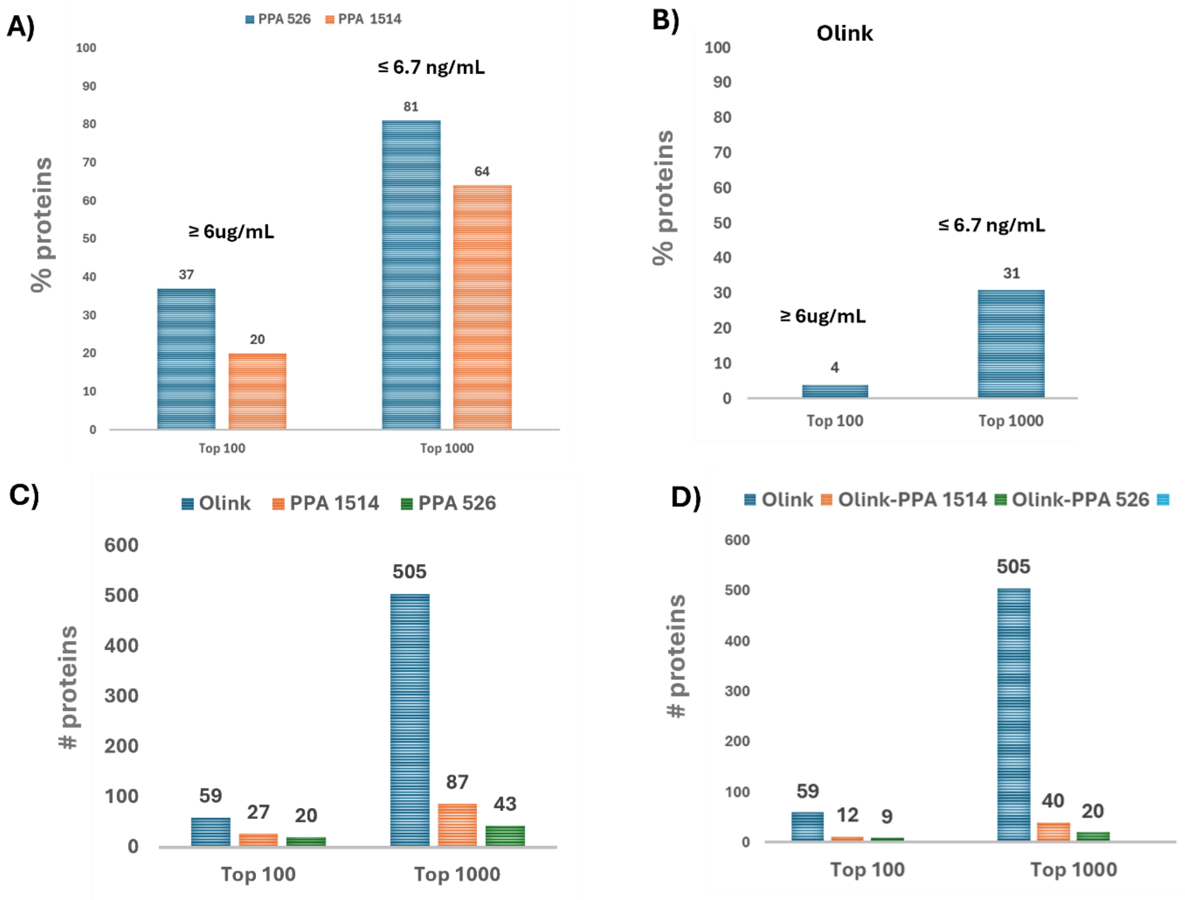

**Figure S11. Abundance-based classification of plasma protein panels using HPA mass spectrometry (BUP) quantification. A)** Percentage of proteins from the **PPA 1514** and **PPA 526** panels that rank among the Top 100 or Top 1000 most abundant plasma proteins in the Human Protein Atlas (HPA). **B)** Percentage of proteins from the Olink HT 3K panel that ranks among the Top 100 or Top 1000 most abundant plasma proteins in the HPA. **C)** Number of proteins classified as Top 100 or Top 1000 across all panels. **D)** Number of proteins classified as Top 100 or Top 1000 in the Olink 3K panel, and numbers of proteins shared between Olink 3K and **PPA 1514** or **PPA 526** panels. Top 100 and Top 1000 proteins are defined according to their abundance rank in the HPA. Proteins in the Top 100 and Top 1000 categories have plasma concentrations of ≤6 µg/mL and ≤6.7 ng/mL, respectively, as estimated by HPA and BUP mass spectrometry.

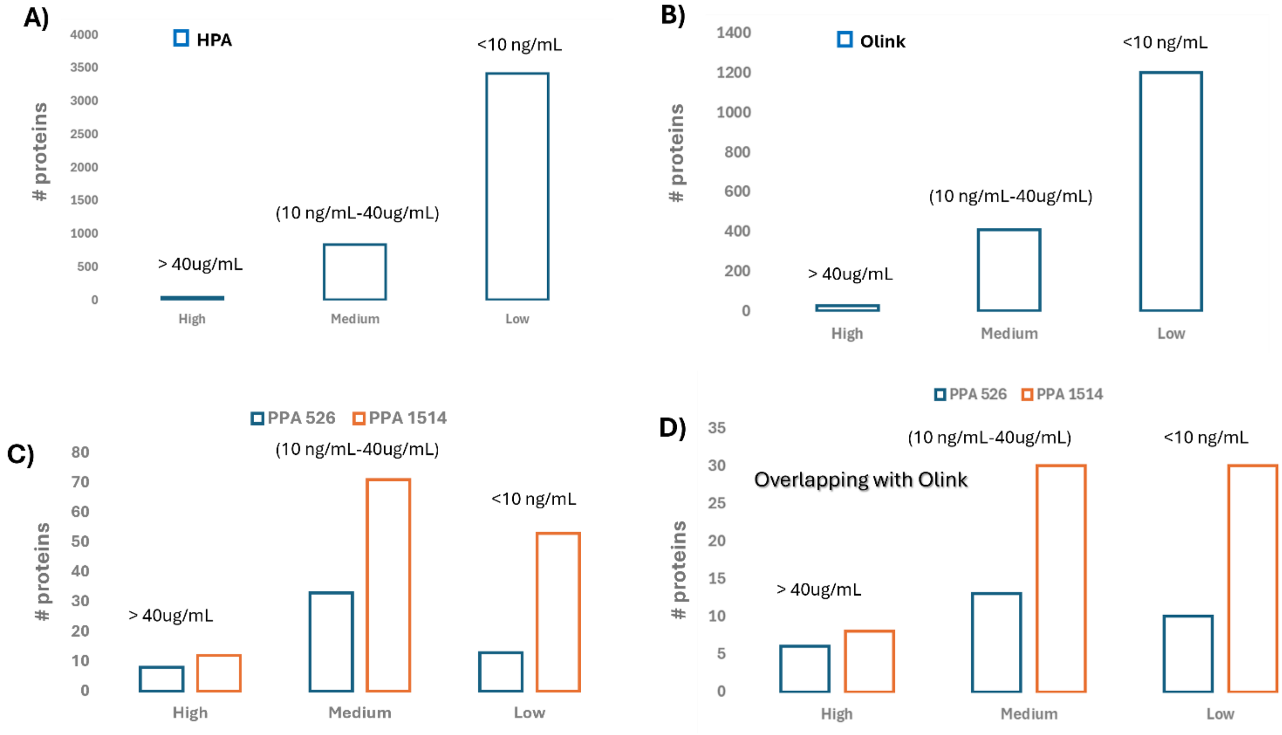

**Figure S12. Abundance based classification of plasma proteins using reported concentration values from the Human Protein Atlas (HPA), grouped into low, medium, and high abundance categories. A)** All plasma proteins are reported in the HPA database. **B)** Proteins included in the Olink 3K panel. **C)** Proteins included in **PPA 1514** and **PPA 526** panels. **D)** Proteins shared between the Olink 3K panel and the **PPA 1514** or **PPA 526** panels. Protein abundance categories are defined based on HPA BUP mass spectrometry estimates as follows: high abundance (>40 µg/mL), medium abundance (10 ng/mL to 40 µg/mL), and low abundance (<10 ng/mL).

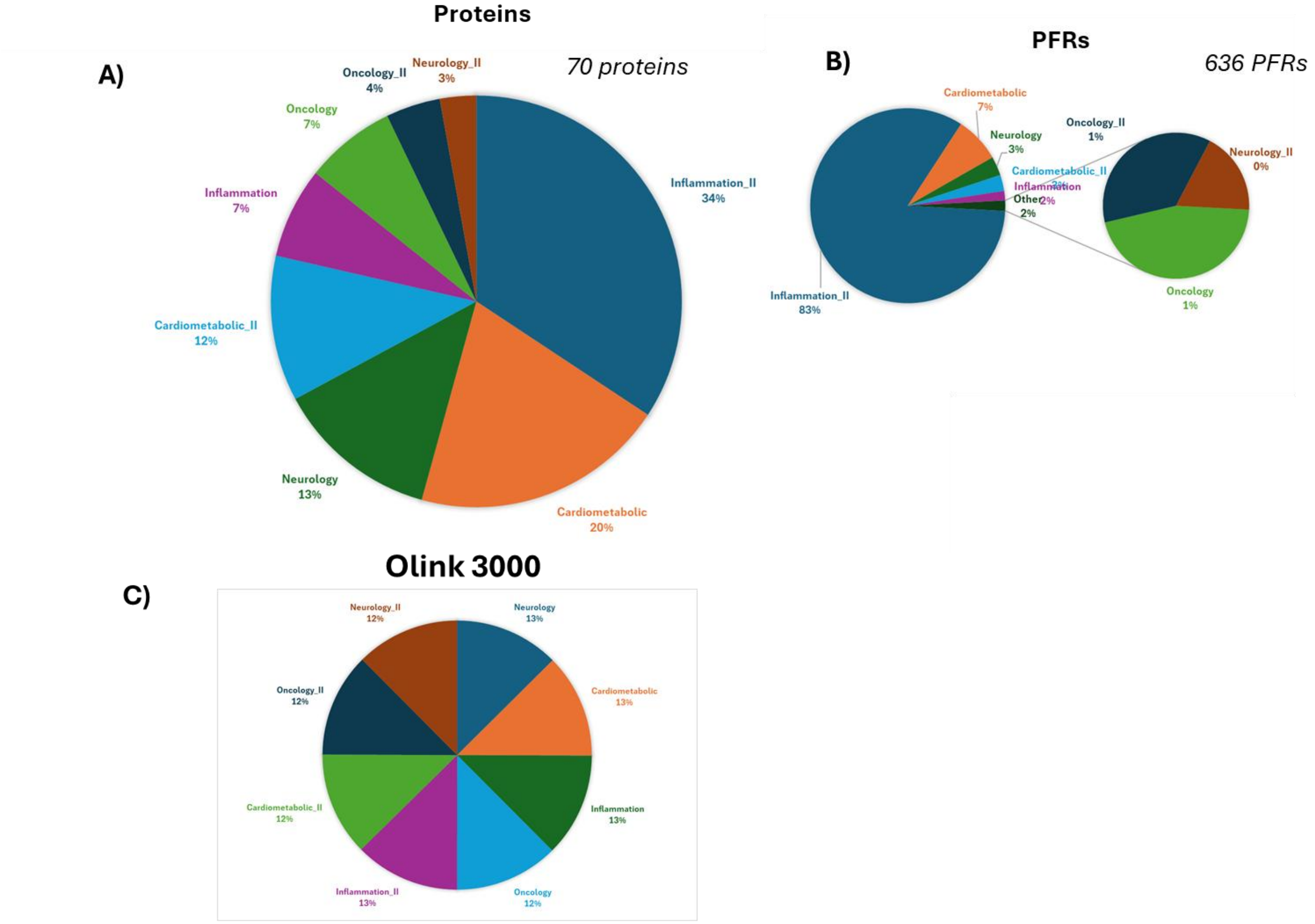

**Figure S13. Functional distribution of plasma proteins shared between Olink 3K and PPA 1514 panels. A)** Pie chart showing the distribution of functional categories among the 70 proteins shared between the **PPA 1514** and Olink 3K panels. **B)** Pie chart showing the distribution of proteoform (PFRs) derived from the 70 shared proteins shown in panel A of the figure. **C)** Pie chart showing the functional category distribution of all proteins included in the Olink 3K panel. Functional categories are based on annotations provided by Olink in their 3K HT panel.

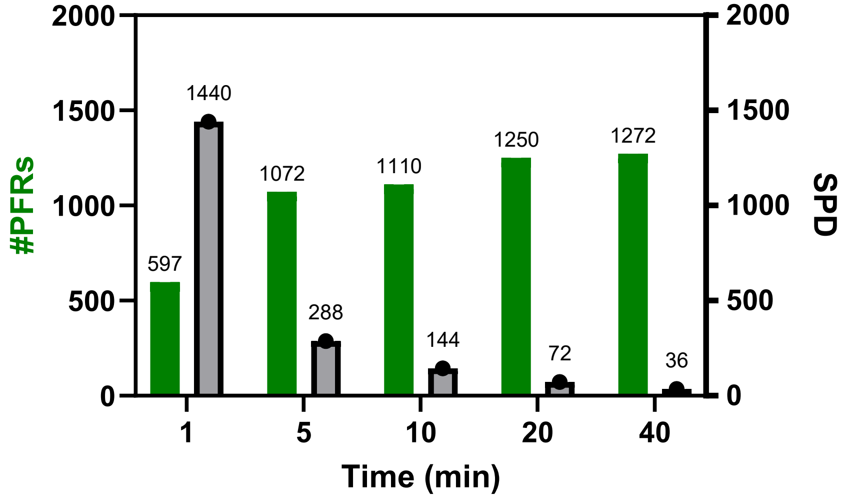

**Figure S14. Estimation of proteoform number and sample throughput possible for PPA 1514 FAIMS-enabled plasma analyses.** Green bars (left axis) represent the number of proteoform features (PFRs) identifiable per run at each acquisition time (x-axis), while gray bars (right axis) indicate the expected number of samples per day (SPD) achievable, estimated assuming continuous acquisition (1440 min $day^{-1}$). Numeric values above bars denote the corresponding PFR and SPD estimates.

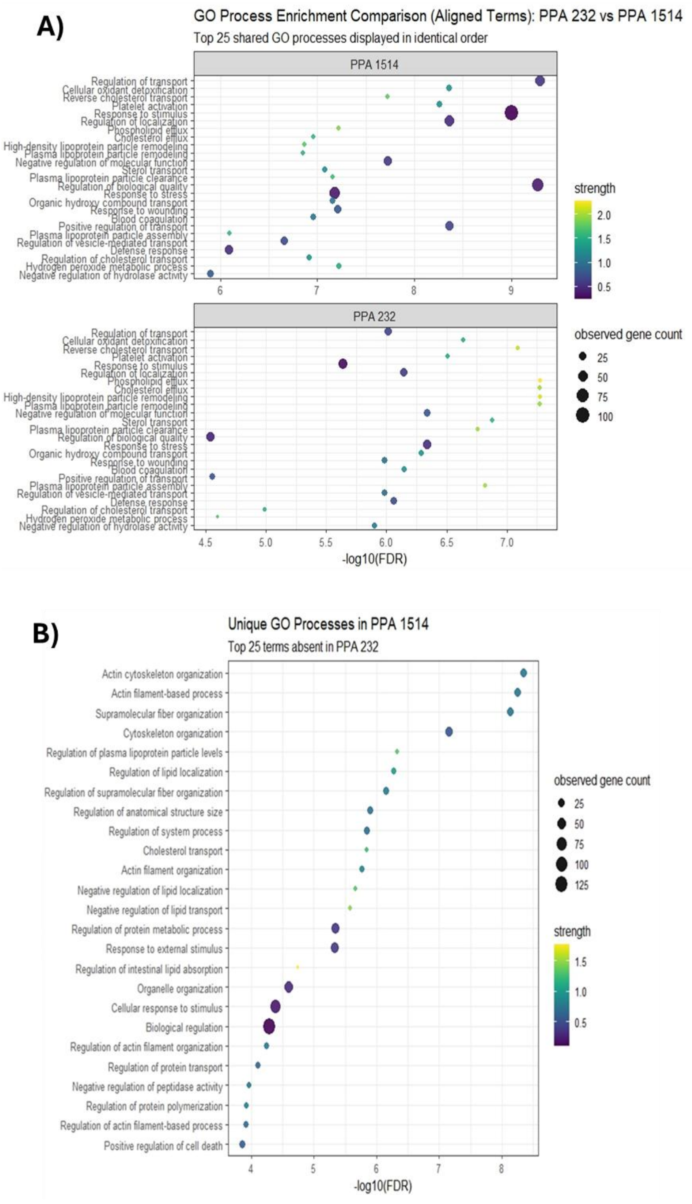

**Figure S15. Comparative analysis of biological processes enriched when including PPA 232 which uses the simple "dilute and shoot" approach. A)** Functional Gene Ontology (GO) biological process enrichment analysis highlighting the top shared processes enriched by **PPA 232** and **PPA 1514. B)** Biological processes uniquely enriched by **PPA 1514** compared with **PPA 232.**

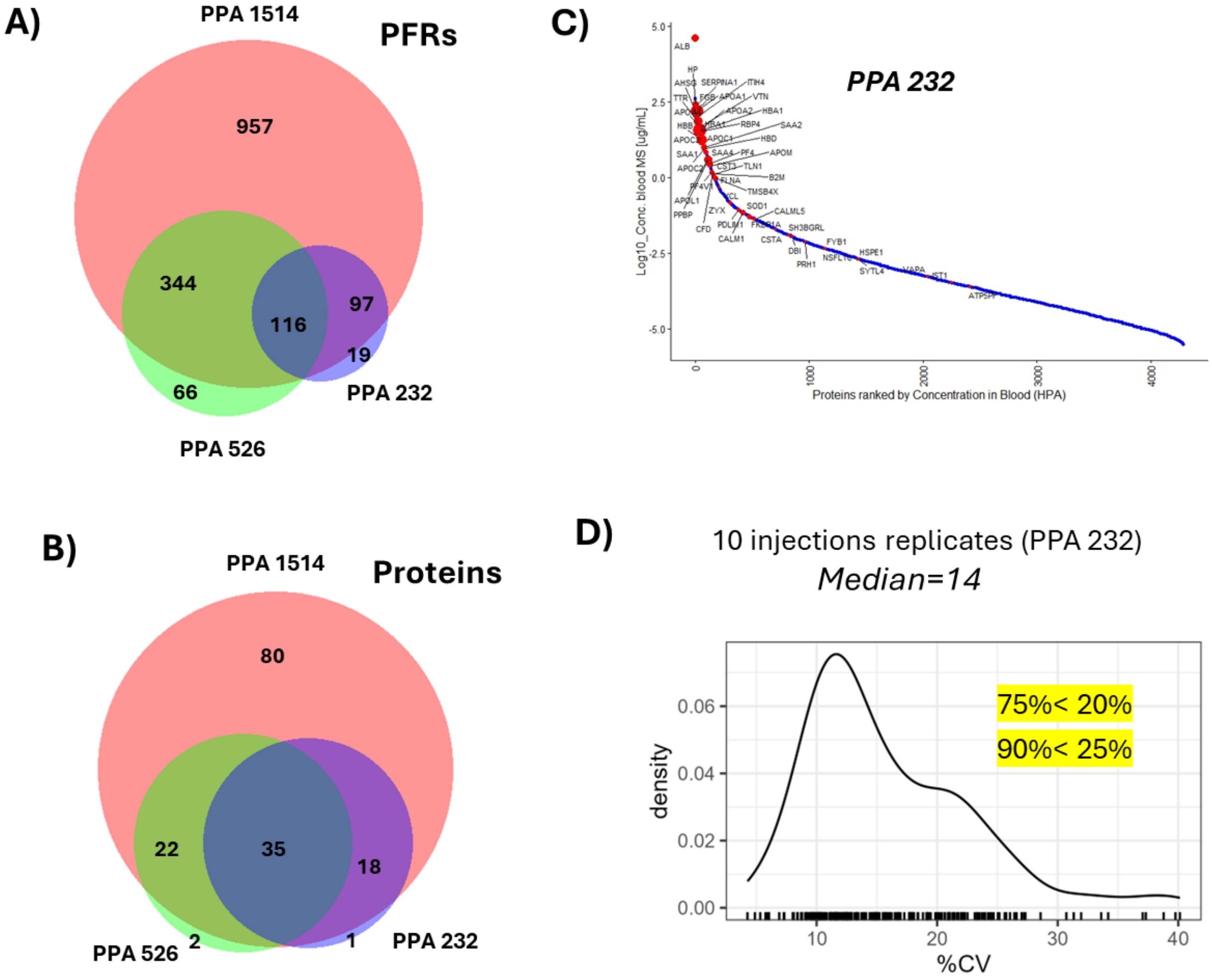

**Figure S16. Comparative analysis of proteoform panels, including PPA 232 which uses the simple "dilute and shoot" approach.** Comparative analysis of proteoform panels, including **PPA 232** which uses the simple "dilute and shoot" approach. **A)** Venn diagram showing overlap among proteoforms identified using **PPA 1514**, **PPA 526**, and **PPA 232**. **B)** Venn diagram depicting an overlap of proteins identified by the PPAs described in (A). **C)** Waterfall plots illustrating the dynamic range of proteins identified using **PPA 232**. **D)** Coefficients of variation (CVs) of proteoform ion counts calculated from 10 technical I²MS injections of a reference sample using **PPA 232**.
